# Cell-scale gene-expression measurements in *Vibrio cholerae* biofilms reveal spatiotemporal patterns underlying development

**DOI:** 10.1101/2024.07.17.603784

**Authors:** Grace E. Johnson, Chenyi Fei, Ned S. Wingreen, Bonnie L. Bassler

**Author notes:** Present address: Department of Mathematics, Massachusetts Institute of Technology, Cambridge, MA 02139.

## Abstract

Bacteria commonly exist in multicellular, surface-attached communities called biofilms. Biofilms are central to ecology, medicine, and industry. The *Vibrio cholerae* pathogen forms biofilms from single founder cells that, via cell division, mature into three-dimensional structures with distinct, yet reproducible, regional architectures. To define mechanisms underlying biofilm developmental transitions, we establish a single-molecule fluorescence in situ hybridization (smFISH) approach that enables accurate quantitation of spatiotemporal gene-expression patterns in biofilms at cell-scale resolution. smFISH analyses of *V. cholerae* biofilm regulatory and structural genes demonstrate that, as biofilms mature, overall matrix gene expression decreases, and simultaneously, a pattern emerges in which matrix gene expression becomes largely confined to peripheral biofilm cells. Both quorum sensing and c-di-GMP-signaling are required to generate the proper temporal pattern of matrix gene expression. Quorum sensing autoinducer levels are uniform across the biofilm, and thus, c-di-GMP-signaling alone sets the regional matrix gene expression pattern. The smFISH strategy provides insight into mechanisms conferring particular fates to individual biofilm cells.

## Introduction

Bacteria commonly exist in spatially-organized, surface-attached communities called biofilms. Biofilms provide advantages to the cells residing within them, including protection from environmental insults such as antimicrobials, predators, and mechanical perturbation [1–4]. Critical to the biofilm lifestyle is the production of an extracellular matrix that attaches cells to the surface and to one another [5]. Biofilm cells can degrade the matrix, exit the community, and return to the individual, planktonic lifestyle in a process called dispersal [6,7]. The ability to transition between biofilm and planktonic lifestyles allows bacteria to respond to changing environments and, moreover, is required for many pathogens to successfully cause disease [6]. For example, *Vibrio cholerae*, the etiological agent of the disease cholera and the model organism used in this work, forms biofilms both in its marine niche and in human hosts, and successive rounds of biofilm formation and dispersal are key to transmission of cholera disease [8,9].

In *V. cholerae*, the major component of the matrix is an exopolysaccharide called *Vibrio* polysaccharide (VPS) [10]. The enzymes necessary for VPS synthesis are encoded in two operons, *vpsI* and *vpsII*. In addition to VPS, the *V. cholerae* extracellular matrix contains three major proteins: RbmA, which promotes cell-cell adhesion, Bap1, important for cell-surface attachment, and RbmC, which, along with Bap1 and VPS, forms envelopes that encapsulate cell clusters [11–13]. Expression of the two *vps* operons, *rbmA*, *bap1*, and *rbmC*, and correspondingly the production of matrix, is regulated by two sensory inputs, the small second messenger molecule cyclic diguanylate (c-di-GMP) and quorum-sensing-mediated chemical communication (QS) [14–16].

c-di-GMP controls *V. cholerae* matrix production by binding to and activating two transcription factors, VpsR and VpsT, which in turn, drive transcription of matrix genes (Fig 1) [15]. c-di-GMP is synthesized and degraded by enzymes with diguanylate cyclase (DGC) and phosphodiesterase (PDE) activities, respectively [17]. *V. cholerae* DGC and PDE activities are controlled by environmental stimuli. One such stimulus is the polyamine norspermidine (Nspd). Nspd, via its NspS receptor, drives increases in c-di-GMP production by interacting with the bifunctional enzyme MbaA, activating its DGC activity and suppressing its PDE activity (Fig 1) [18]. *V. cholerae* does not secrete detectable Nspd under laboratory-growth conditions. Thus, exogenous administration of Nspd can be used to modulate cytoplasmic c-di-GMP levels, as described previously [19]. We employ this strategy in the present work.

**Fig 1.**
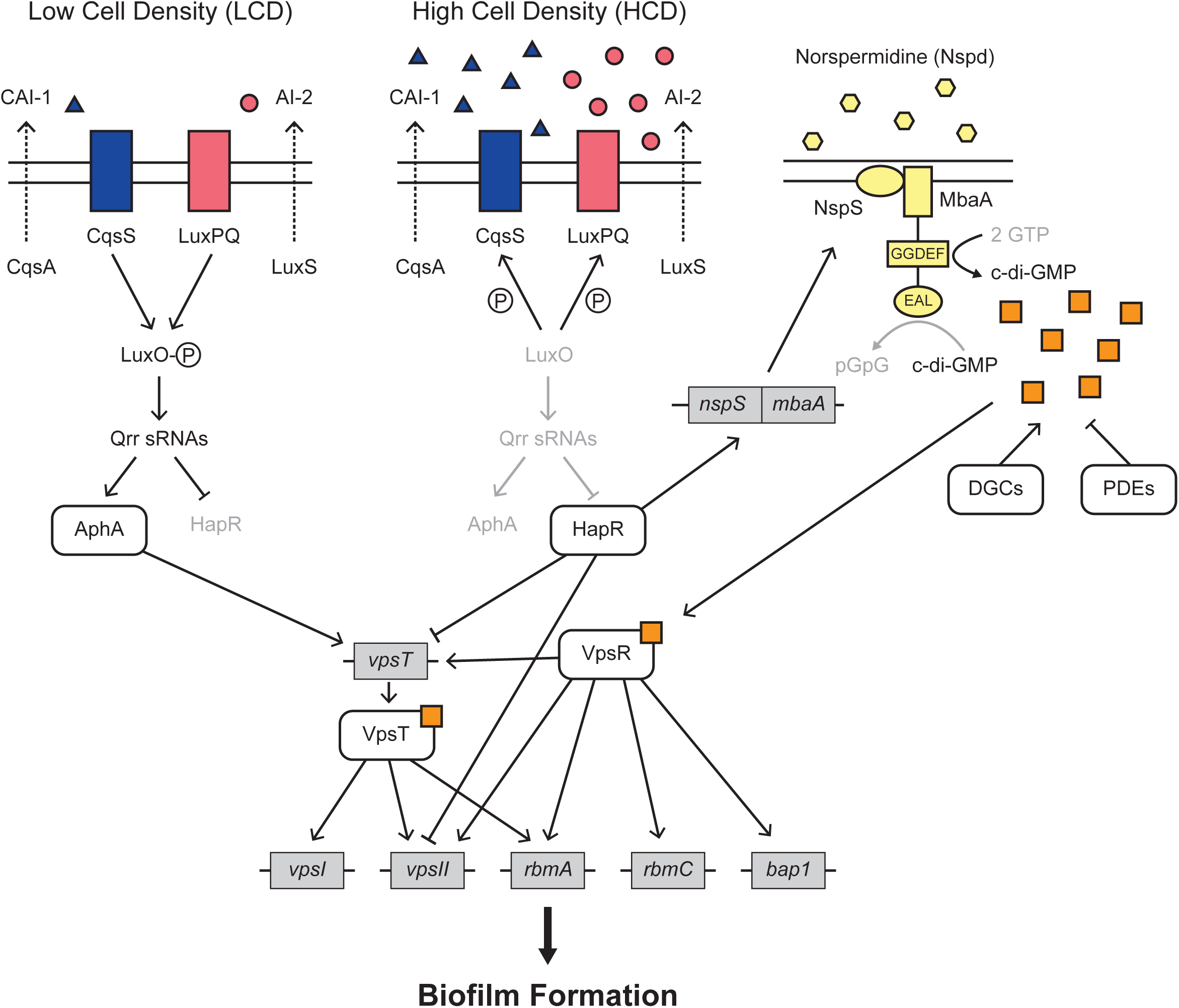
Regulation of biofilm formation in *V. cholerae*. Simplified model of regulation of biofilm formation by quorum sensing (QS), c-di-GMP signaling, and norspermidine (Nspd). See text for full details. Blue triangles represent CAI-1; pink circles represent AI-2; yellow hexagons represent Nspd; orange squares represent c-di-GMP. The P in a circle represents phosphate. The GGDEF and EAL designations represent the motifs on MbaA that confer the DGC and PDE activities, respectively.

QS is the process of bacterial communication in which cells produce, release, and detect extracellular signal molecules called autoinducers to regulate group behaviors, including biofilm formation (Fig 1) [20,21]. *V. cholerae* possesses multiple QS autoinducer-receptor pairs, two of which are relevant to the present work. The first autoinducer is *cholerae* autoinducer-1 (CAI-1; (*S*)-3-hydroxytridecan-4-one), which is synthesized by CqsA and detected by the CqsS receptor [22,23]. The second autoinducer is autoinducer-2 (AI-2; (2S,4*S*)-2-methyl-2,3,3,4-tetrahydroxytetrahydrofuran borate), which is synthesized by LuxS and detected by the LuxPQ receptor [24,25]. At low cell densities (LCD), when autoinducer concentrations are low, the unbound CqsS and LuxPQ receptors act as kinases and shuttle phosphate to the LuxO response regulator [26,27]. LuxO∼P activates expression of genes encoding a set of small RNAs (sRNAs) called the Qrr sRNAs, which activate production of the master LCD regulator AphA and repress production of the master high-cell-density (HCD) regulator HapR [28,29]. Conversely, at HCD, when CAI-1 and AI-2 have accumulated, the autoinducer-bound receptors act as phosphatases, LuxO is dephosphorylated, and transcription of the *qrr* genes terminates [26–28]. As a result, AphA production ceases, and HapR production commences [29]. HapR represses transcription of *vpsT* and the *vpsII* operon [30,31]. Thus, at LCD, when HapR levels are low, matrix production, and consequently, biofilm formation, occur, while at HCD, when HapR levels are high, matrix production is repressed, promoting biofilm disassembly and cell dispersal (Fig 1) [32].

Analogous to eukaryotic embryos, bacterial biofilms often arise from single cells that develop into multicellular structures via cell division [33]. Recent advances in microscopy have made it possible to image and track individual live bacteria over the initial phase of biofilm development, revealing that cells reproducibly adopt unique cell fates depending on their locations in the emerging biofilm [33–35]. We know that cells are organized into distinct clusters surrounded by non-uniform distributions of matrix components [12]. Furthermore, individual-cell characteristics, such as orientation, packing with other cells, and trajectories from birth to final destination have been quantified and exhibit location-dependent differences [34,35]. Nonetheless, a mechanistic understanding of how individual cell fates arise is still lacking because accurately measuring individual-cell gene-expression patterns in space and time in developing biofilms has remained a hurdle.

Here, we describe the development and application of a biofilm-specific single-molecule fluorescence in situ hybridization (smFISH) approach that overcomes many of the challenges that have previously limited individual cell gene-expression measurements in biofilms. We first deliver the biofilm-specific smFISH strategy and validate that it accurately quantifies cell-scale gene expression in biofilms. Second, we use the smFISH approach to quantify the spatiotemporal expression patterns of biofilm regulators and downstream structural components. We find that QS is required to establish the overall temporal expression pattern of key biofilm matrix genes, while c-di-GMP signaling confines expression of those matrix genes to distinct biofilm sub-regions. These cell-scale gene-expression measurements provide insight into the molecular mechanisms that confer particular fates to individual cells in *V. cholerae* biofilms.

## Results

### Deploying smFISH for accurate quantitation of cell-scale gene expression in bacterial biofilms

Understanding how individual bacteria residing in biofilm communities adopt location-specific cell fates requires measurements of spatiotemporal gene expression at cell-scale resolution. Traditional fluorescent reporters are inadequate for this task because their output is dampened by the low oxygen levels that exist in biofilms, which prohibits proper reporter maturation [36]. Indeed, in *V. cholerae* biofilms, signal from multiple fluorophores rapidly declines over time, well before biofilms reach maturity (S1A Fig), making fluorescent reporters poor proxies of gene expression in this context. One solution to this problem employed recently is to grow biofilms under flow, and thus supply a constant source of oxygen [37,38]. While this strategy eliminates fluorescent signal decay, it prevents the natural accumulation of small signal molecules, for example QS autoinducers, that are required to properly regulate the biofilm lifecycle.

**S1 Fig. Confocal microscopy biases regarding constitutive fluorescent reporters and stains** (A) Maximum projection confocal microscopy images of mScarlet and mNG fluorescence over time in a representative *V. cholerae* biofilm harboring pTac-*mScarlet* and pTac-*mNG*. (B) Signal from DAPI staining in the first in-focus *z* slice and *xz* and *yz* cross sections of a mature *V. cholerae* biofilm. All scale bars represent 5 µm.

Compounding issues with traditional fluorescent reporters, biases inherent to confocal imaging preclude accurate quantitation of spatial fluorescence patterns in biofilms [35]. *z*-positional bias exists because captured fluorescence signal declines as a function of distance from the objective. Radial (*r*)-positional bias is likely caused by the biofilm’s dome shape. Specifically, the center of the biofilm is thicker and contains more layers of cells than does the edge, so background fluorescence from out-of-focus *z* planes is higher at the biofilm center than at the periphery. Indeed, thinner biofilms with fewer *z* planes display less dramatic *r*-positional bias. Consistent with these imaging biases, we show that signal from DAPI stained biofilms, which is expected to be spatially uniform, decreases from the bottom to the top and from the inside to the outside of *V. cholerae* biofilms (S1B Fig).

With the aim of overcoming the above limitations and developing a robust tool for measuring cell-scale gene expression in bacterial biofilms, we employed smFISH technology. smFISH has been successfully used in planktonic bacteria to accurately measure single-cell gene expression at ranges from <1 to 100 mRNA molecules per cell [39,40]. smFISH does not require an oxygen-dependent maturation step, making it suitable for use in bacterial biofilms. Indeed, sequential smFISH (seqFISH) was recently used to measure the expression of over 100 genes in individual *Pseudomonas aeruginosa* biofilm cells [41]. Below, we combine smFISH with confocal imaging and a biofilm-specific downstream analysis pipeline to accurately quantify cell-scale gene expression over space and time in *V. cholerae* biofilms. Our approach provides a powerful tool for such analyses, including for transcripts that display only low-level expression and for proteins that are recalcitrant to tagging.

Our first goal was to assess the imaging artifacts associated with our smFISH strategy independent of any biological gene-expression patterns. To do this, we measured the smFISH fluorescence signal from a gene expressed uniformly across the biofilm. We grew *V. cholerae* biofilms containing arabinose-inducible *m-neon green (mNG*) integrated into the chromosome. Previous reports demonstrated uniform cell growth across the biofilm under our conditions [35], and thus, *mNG* expression levels should track with the exogenously supplied arabinose inducer concentration. To ensure equivalent arabinose penetration throughout the biofilm, it was added at the start of the experiment, prior to biofilm formation. Importantly, while *V. cholerae* cells are able to import arabinose, they are not known to metabolize it, and thus arabinose concentrations should remain spatially uniform and constant over the course of growth [42]. We grew biofilms in the presence of high and low concentrations of arabinose and measured *mNG* transcripts with smFISH probes labeled with each of the three fluorescent dyes used throughout the remainder of this work. The fluorescence signal from the mNG protein does not survive the fixation and smFISH hybridization process, and thus does not add to nor interfere with quantitation of smFISH fluorescence signal. This strategy allowed us to ascertain fluorescence signal biases across entire biofilms for each fluorescent probe, and over a range of transcript levels. *mNG* expression levels assessed by each fluorophore in biofilms paralleled those in planktonic cells grown with the corresponding inducer concentration (S2A Fig).

**S2 Fig. Quantitation of spatial per-cell-volume smFISH fluorescence signal.** (A) Average per-cell-volume *mNG* smFISH fluorescence signal measured in *V. cholerae* planktonic and biofilm cells. 0.2% or 0.0375% arabinose was provided, as indicated by shaded blue boxes denoted High and Low Expression, respectively. 100 µM Nspd was included in all cases. *mNG* expression was measured by smFISH using probes labeled with one of three fluorophores, as indicated. Values for biofilm cells represent the mean normalized per-cell-volume fluorescence signal, calculated as described in (B), across *n* = 10-12 biofilms. The small, medium, and large circular symbols represent biofilms with approximate biovolumes of 2^9^, 2^11^, and 2^14^ µm^3^, respectively. Error bars denote standard deviations, which are in some cases smaller than the sizes of the symbols used in the plots. Values for planktonic cells represent the average across all cells in single replicate experiments; error bars are excluded. (B) Schematic overview of quantitation of per-cell-volume fluorescence signal. Briefly, using BiofilmQ [43], biofilms are broken into cubes with side lengths of 2.32 µm and cells within these cubes are identified using the DAPI channel. The cell volume (*V_c_*) for each cube is calculated as the fraction of the total cube volume occupied by cell mass. Puncta from smFISH fluorescence signal are detected as local maxima, and the integrated, background-subtracted punctum intensity calculated (*I*). The per-cell-volume fluorescence signal is subsequently calculated as the ratio of the puncta intensity and the cell volume (*I*/*V_c_*). For full details see Methods: smFISH image analysis. (C) Probability density curves of the distributions of individual smFISH puncta intensities for the *mNG* transcript measured under the Low Expression condition, *hapR* measured at LCD, and *qrr*4 and *vpsL* measured at HCD. The peaks of the probability curves were used to estimate the intensities of a single mRNA molecule for each transcript. The minimum (0.5) and maximum (1.7) puncta intensities are highlighted with vertical lines. (D) Schematic overview of quantitation of spatial gene expression patterns in biofilms. Briefly, cells are grouped by their *r* and *z* positions, and the average per-cell-volume fluorescence signal in each group is calculated. For full details, see Methods: Analysis of spatial gene-expression patterns at cell-scale resolution.

We quantified smFISH fluorescence signal at cell-scale resolution (2.32 µm) by dividing biofilms into cell-sized cubes [43] (here forward referred to interchangeably as “cell cubes” or “cells” for simplicity) with 2.32 µm sides and calculating the volume-normalized smFISH signal in each cell cube (see S2B Fig, Methods: smFISH image analysis). We refer to resulting values henceforth as ‘per-cell-volume fluorescence signal’. Of note, per-cell-volume fluorescence signal is reported here and throughout this work in arbitrary units (AUs). These values can be converted to transcript numbers by identifying the fluorescence distributions of individual puncta reporting on transcripts that are produced at low levels, where the signal from most puncta is predicted to arise from an individual mRNA molecule (S2C Fig and [39]). In our experiments, individual mRNA puncta intensities cluster around 1 and differ by no more than 2-fold across fluorophores and genes. Thus, 1 AU corresponds to approximately 0.5 – 2 mRNA transcripts. To quantify spatial smFISH fluorescence signal patterns, we grouped cells across replicate biofilms by their *r* and *z* positions.

We calculated the average per-cell-volume fluorescence signal for each group of cells relative to the average per-cell-volume fluorescence signal of a group of cells located near the center of the biofilm (S2D Fig, Methods: Analysis of spatial gene-expression patterns at cell-scale resolution). The *r* and *z* positions of each group, and the corresponding average per-cell-volume fluorescence signals relative to that in the reference group of cells with position *r, z* = 2.32, 2.32 µm, can be visualized as a heatmap, as in Fig 2A.

**Fig 2.**
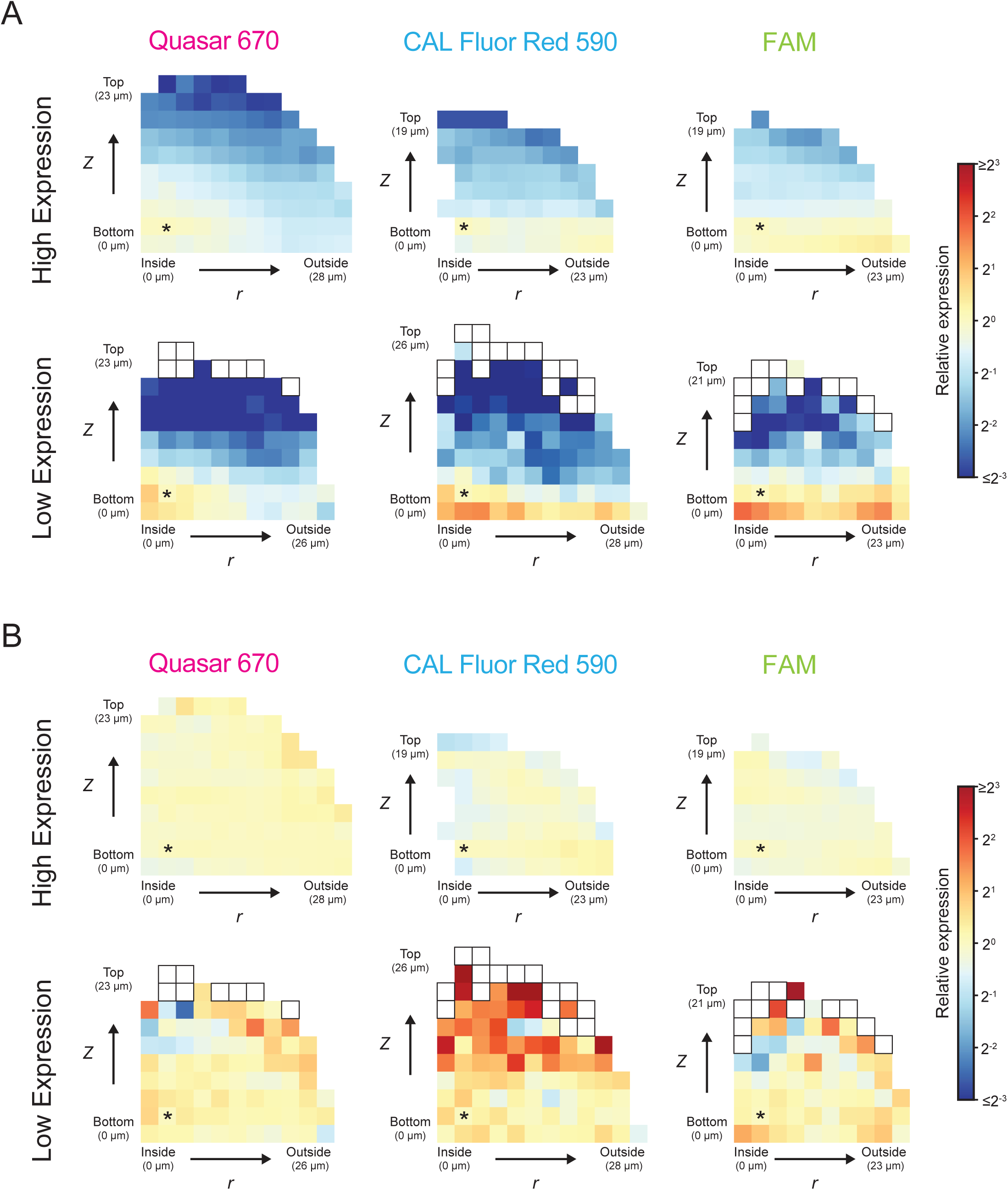
Quantitation of cell-scale gene expression in *V. cholerae* biofilms. (A) Heatmaps showing relative, non-corrected spatial *mNG* per-cell-volume smFISH fluorescence signal data as a function of position *r, z* in biofilms of *V. cholerae* carrying pBad-*mNG*. Biofilms were grown in the presence of 0.2% or 0.0375% arabinose, denoted High and Low Expression, respectively. 100 µM Nspd was included in all cases. *mNG* expression was measured by smFISH using probes labeled with one of three fluorophores, as indicated. Cells are grouped into bins with size Δ*r,* Δ*z* = 2.32, 2.32 µm. Gene expression in each bin is represented as a relative value, where the average per-cell-volume fluorescence signal across all cells in a bin is normalized to the average per-cell-volume fluorescence signal across all cells in the bin denoted with the asterisk (bin *r, z* = 2.32, 2.32 µm). White boxes represent zero values. (B) Heatmaps as in (A) showing the relative, corrected per-cell-volume fluorescence signal values. All data represent *n* = 10-12 biofilms.

Absent fluorescence signal biases, and given uniform expression of *mNG*, one would expect unvarying fluorescence signal across all cells in the biofilm. This is not the case. Rather, spatially-dependent fluorescence signal differences are observed, in which average per-cell-volume fluorescence signal is highest in cells closest to the center of the biofilm, and lowest in cells most distant from the center (Fig 2A, S3A Fig). The pattern is consistent with the known *z* and *r* positional biases of confocal imaging described above. Crucially, the fluorescence signal pattern is the opposite of what would be predicted if the arabinose inducer and/or *mNG* probes could not penetrate the biofilms. In both of those cases, one would expect lower per-cell-volume fluorescence signal in the center of the biofilm and higher per-cell-volume fluorescence signal at the biofilm periphery. While we cannot eliminate the possibility that the obtained fluorescence pattern arises from higher arabinose concentrations in the biofilm center than at the periphery, arabinose accumulation at the biofilm core is unlikely given that *V. cholerae* does not metabolize arabinose. We know of no other *a priori* reason for why arabinose concentration would differ across the biofilm. Thus, the spatial fluorescence signal pattern observed here most likely stems from constitutive gene expression and arises entirely from the expected confocal imaging artifacts.

**S3 Fig. Validation of fluorescent signal quantitation in *V. cholerae* biofilms.** (A) Average per-cell-volume fluorescence signal versus distance to the biofilm center (*r, z* = 0, 3.5 µm) is shown for each fluorophore as indicated. The corrected and non-corrected data are shown for biofilms with high and low *mNG* expression. All data represent *n* = 10-12 biofilms. (B) Heatmaps as in Fig 2A of the main text showing the relative corrected data for the three fluorophores for biofilms of additional sizes. Data represent *n* = 10-12 replicate biofilms. (C) As in (A), for the data shown in (B). Black symbols represent biofilms in the left column (smaller biofilms) and white symbols represent biofilms in the right column (larger biofilms) of (B).

To account for these artifacts, we developed a mathematical model to quantify the above fluorescence signal positional biases for each fluorophore and across different gene-expression levels. When we use this method to correct the *mNG* expression data, we observe uniform average *mNG* per-cell-volume fluorescence signal throughout biofilms, consistent with our expectation for a constitutively expressed gene (Fig 2B, S3A Fig). The average per-cell-volume fluorescence signal appears less uniform in biofilms that display low *mNG* expression as a consequence of higher expression variation between cells. The average per-cell-volume fluorescence signal is most noisy in bins far from the surface, the region with the least fluorescence signal. However, 97% of bins across biofilms with low *mNG* expression have a *z*-score between -0.25 and 0.25, i.e., they deviate from the average per-cell-volume fluorescence signal of the center bin by no more than one quarter of a standard deviation, demonstrating that there are minimal fluorescence signal output differences between groups of cells. The model is not only able to correct spatial fluorescence signal biases in the large biofilms used to generate the model, but can also be directly applied without modification to accurately correct biases from smaller biofilms grown with identical concentrations of arabinose inducer (S3B, C Fig).

To assess the applicability of our model, we measured the expression of two constitutively expressed housekeeping genes, *gyrA,* encoding DNA gyrase subunit A, and *polA,* encoding DNA polymerase I, using smFISH. We observe consistent expression of both genes over the course of biofilm development (S4A Fig) at levels comparable to those previously reported (the ratios of *gyrA*:*polA* expression measured by smFISH here and measured in planktonic *V. cholerae* cells by RNA-seq in [44] are 3.5 and 2.8, respectively). Following correction using our model, we measure uniform expression of *gyrA* and *polA*, with expression varying less than 2-fold across all biofilm positions (S4B-E Fig). These data highlight the power of our model to correct for imaging biases irrespective of the specific transcript being measured, and they demonstrate how the correction can be used to obtain accurate cell-scale spatial gene-expression patterns in biofilms for, in principle, any gene of interest. These data also support our assumption that confocal imaging biases, and not regional differences in arabinose concentration, underlie the spatial pattern used to generate our model. If non-uniform arabinose-driven *mNG* expression occurred in biofilms, our model would not accurately correct the spatial expression profiles of other uniformly expressed genes.

**S4 Fig. Validation of spatial correction model.** (A) Average per-cell-volume *gyrA* and *polA* smFISH fluorescence signal before (top) and after (bottom) correction using the model described in the text (See Methods: Spatial correction model) across replicate *V. cholerae* biofilms grown with 100 µM Nspd as a function of the average biofilm biovolume. Dotted lines denote the average smFISH fluorescence signal across all biovolumes. Error bars denote standard deviations across *n* = 9-10 replicate biofilms, which are in some cases smaller than the sizes of the symbols used in the plots. (B) (Top) Representative confocal microscopy images showing DAPI, and *gyrA,* and *polA* smFISH fluorescence signal in biofilms with the largest biovolumes shown in (A). Images represent maximum projections of the first four in-focus *z* slices. Scale bars represent 5 µm. (Bottom) Heatmaps showing average *gyrA* and *polA* non-corrected (upper) and corrected (lower) per-cell-volume smFISH fluorescence signal across *n* = 10 replicate biofilms as a function of *x, y* position (with replicate biofilms aligned around their center positions) for biofilm cells with *z* positions between 0 and 4.68 µm in biofilms with the largest biovolumes shown in (A). Scale bars represent 5 µm. (C) Non-corrected (top) and corrected (bottom) relative average per-cell-volume fluorescence signal as a function of distance from the periphery of the biofilm for the data shown in (B). Values are relative to the average per-cell-volume fluorescence signal of cells located at a distance of 2 µm from the biofilm periphery for each gene. (D) Heatmaps of *gyrA* and *polA* non-corrected (top) and corrected (bottom) average per-cell-volume fluorescence signal relative to the average per-cell-volume fluorescence signal in the bins denoted with asterisks are shown as in Fig 2B for *n* = 10 replicate biofilms with the largest biovolumes shown in (A). (E) Average non-corrected (top) and corrected (bottom) per-cell-volume fluorescence signal versus distance to the biofilm center, as in S3A Fig, for the data shown in (D).

Full details of how we developed the model and how it is applied are provided in Methods. All main text figures show data that has been corrected using our model. All companion plots generated from the non-corrected data are provided in S1 Data, aside from select examples that are included in supplementary figures as noted. Descriptions of per-cell-volume fluorescence signal and expression levels henceforth are also based on corrected data, unless otherwise noted. Using our model to correct for spatial fluorescence signal imaging biases overcomes a major limitation of confocal imaging and provides us with a *V. cholerae* biofilm-specific analysis pipeline that delivers accurate spatial and temporal gene-expression patterns at cell-scale resolution.

### smFISH accurately measures QS gene expression in biofilms

We validated that our smFISH technique could accurately quantify cell-scale gene expression of genes controlling the biofilm lifecycle in *V. cholerae* biofilms across space and time. Here, we focus on the genes encoding the QS LCD and HCD master regulators, *qrr*4 and *hapR*, respectively, and the downstream QS-controlled target gene *vpsL*, the first gene in the *vpsII* operon that encodes VPS biosynthetic enzymes. HapR represses *vpsL* at HCD (Fig 1). Regarding the strategy of using the *qrr*4 gene as the readout for the QS LCD state, the *aphA* gene encoding the LCD master transcription factor would have been the obvious choice (Fig 1). However, AphA is a small protein encoded by a short transcript, and despite multiple attempts, we were unable to design smFISH probes against the *aphA* transcript that yielded measurable fluorescence signal. Attempts to construct AphA fusions to increase transcript length destabilized the mRNAs and they could not be detected by smFISH nor qRT-PCR. The sRNA Qrr4 is also, by definition, a small transcript. Thus, to monitor the QS LCD state, we used smFISH probes against *mNG* encoded under the p*qrr*4 promoter that has previously been demonstrated to accurately report *qrr*4 levels [45,46].

To validate our approach, we compared the fluorescence signal detected by smFISH in biofilms to that from an established technology, smFISH probing planktonic cultures. We know that QS gene expression in *V. cholerae* biofilms can be dramatically different from that in planktonic cells. Thus, for this series of experiments, we exploited a *V. cholerae* strain in which we could control its QS status. Specifically, the *V. cholerae* strain contains only a single QS receptor, LuxPQ, which detects the autoinducer AI-2. The strain also contains a deletion of the gene encoding the AI-2 synthase, *luxS*. Therefore, the level of expression of QS-regulated genes is set exclusively by exogenously supplied AI-2, the amount of which we can precisely control. From here forward, we call this strain ‘the AI-2 sensor strain’. We grew planktonic and biofilm cultures of the AI-2 sensor strain without AI-2 or with a saturating AI-2 concentration (10 µM) [47]. We supplied the AI-2 at the start of the experiment to ensure a constant concentration in the shaking culture and across the biofilm at all developmental stages. This approach enables comparison of gene expression between biofilm and planktonic cells because it forces the strain into the LCD and HCD states, respectively, and in turn drives maximal/minimal *qrr*4 and minimal/maximal *hapR* expression. Moreover, because QS gene expression at each AI-2 concentration is fixed and unvarying in the AI-2 sensor strain, we could assess the accuracy of our measurements over time and space in biofilms. Specifically, we confirmed temporal accuracy by measuring expression in biofilms grown without or with AI-2 over a range of biofilm developmental stages, and we confirmed spatial accuracy by measuring the pattern of gene expression across entire biofilms. Important for our strategy, and mentioned in the Introduction, is that *V. cholerae* forms biofilms at LCD and disperses from biofilms at HCD. Thus, in all cases, the AI-2 sensor strain was grown in the presence of the polyamine norspermidine (denoted Nspd) that prevents biofilm dispersal at HCD [48].

First, regarding the QS regulators *qrr*4 and *hapR*, there is good agreement in the AI-2 sensor strain between expression measured in biofilms and planktonic cells by smFISH (Fig 3A). Specifically, high *qrr*4 expression and low *hapR* expression occur in the -AI-2 (LCD) samples, and low *qrr*4 and high *hapR* expression occur in +AI-2 (HCD) samples (Fig 3A-C). Importantly, the average per-cell-volume expression of *qrr*4 and *hapR* measured both relative to each other and across the LCD and HCD conditions are consistent between planktonic and biofilm cells (Fig 3A, slope of best fit line on a log-log scale for *qrr*4 and *hapR* points is 0.88), confirming the accuracy of our smFISH strategy for measuring gene expression in biofilms. Moreover, for any given condition and gene studied, the global average per-cell-volume fluorescence signal differed by no more than 2-fold over time and across a range of biofilm sizes (Fig 3B). Following application of our spatial correction model to these data (S5A-D Fig), we find consistent and appropriate high or low expression of each gene under each condition across the biofilm (Figs. 3C-E, S5E, S6).

**Fig 3.**
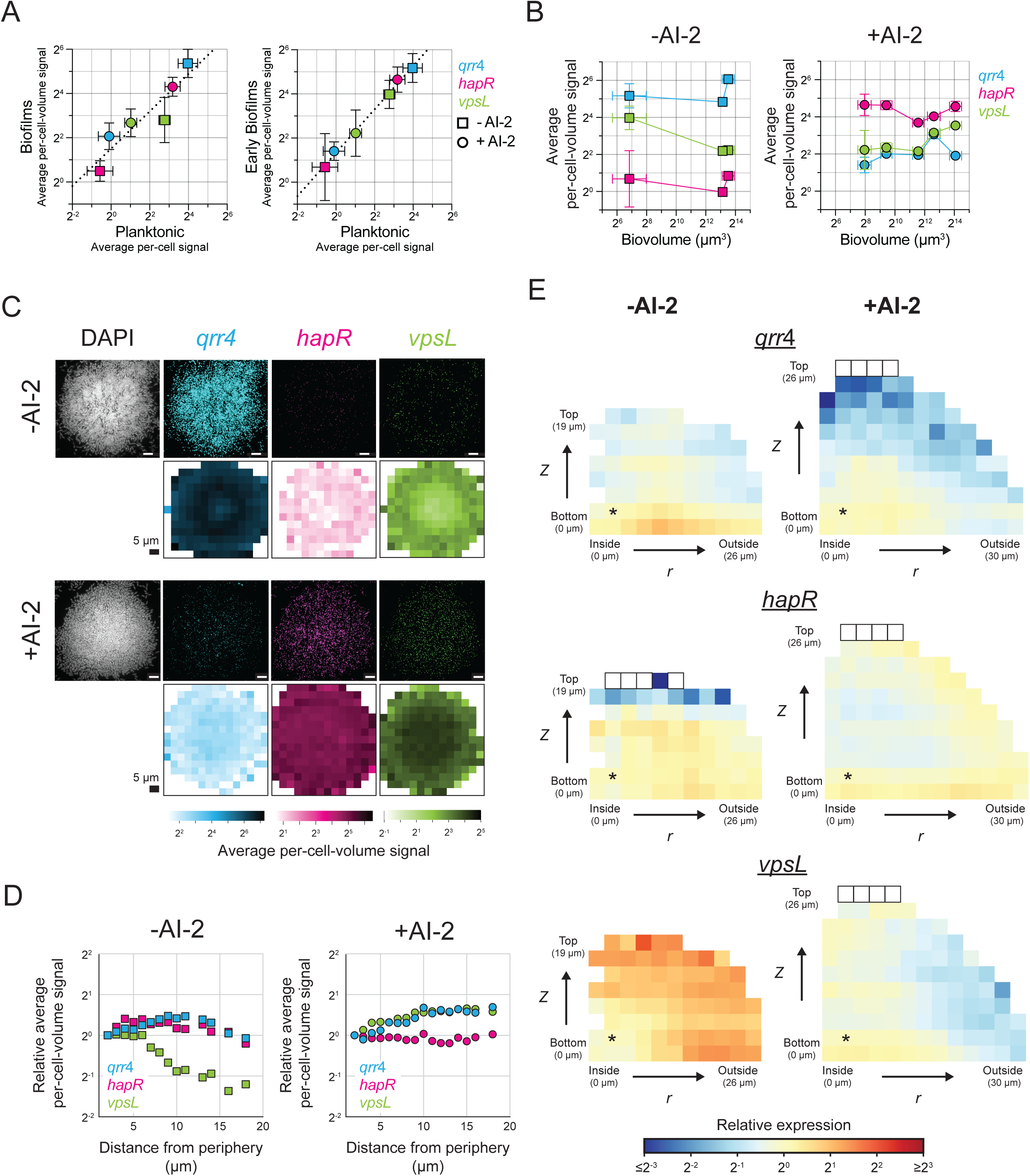
smFISH accurately quantifies QS gene expression in single cells of *V. cholerae* in biofilms. (A) Left panel: Average per-cell-volume *qrr*4 (*mNG*), *hapR,* and *vpsL* smFISH fluorescence signal in *V. cholerae* planktonic and biofilm cells. Values for planktonic samples represent the means of *n* = 3 biological replicates. Values for biofilm samples represent the means across all biofilm sizes, as shown in (B). The slope of the best fit line as shown is 0.80.

The slope of the best fit line excluding *vpsL* is 0.88. Right panel: values for biofilm samples calculated as the means across *n* = 9-10 replicate biofilms with the smallest biovolumes shown in (B). The slope of the best fit line is 0.99. (B) Average per-cell-volume *qrr*4 (*mNG*), *hapR*, and *vpsL* smFISH fluorescence signal across replicate *V. cholerae* biofilms grown without (left) or with (right) 10 µM AI-2 as a function of the average biofilm biovolume. (C) First and third rows: Representative confocal microscopy images showing DAPI, and *qrr*4 (*mNG*), *hapR*, and *vpsL* smFISH fluorescence signals in biofilms with the largest biovolumes shown in (B) grown without (first row) or with (third row) 10 µM AI-2. All biofilms were grown with 100 µM Nspd. Images represent maximum projections of the first four in-focus *z* slices. Scale bars represent 5 µm. Second and fourth rows: Heatmaps showing average *qrr*4 (*mNG*), *hapR*, and *vpsL* per-cell-volume signal across *n* = 10 replicate biofilms as a function of *x, y* position (with replicate biofilms aligned around their center positions) for biofilm cells with *z* positions between 0 and 4.68 µm in biofilms with the largest biovolumes shown in (B) grown without (second row) or with (fourth row) 10 µM AI-2. Scale bars represent 5 µm. (D) Relative average per-cell-volume fluorescence signal as a function of distance from the periphery of the biofilm for the data shown in (C). Values are relative to the average per-cell-volume fluorescence signal of cells located at a distance of 2 µm from the biofilm periphery for each condition and gene. (E) Heatmaps of relative *qrr*4 (*mNG*), *hapR*, and *vpsL* average per-cell-volume fluorescence signal relative to the average per-cell-volume signal in the bin denoted with an asterisk are shown as in Fig 2B for *V. cholerae* biofilms grown without (left) and with 10 µM AI-2 (right), for *n =* 10 replicate biofilms with the largest biovolumes shown in (B). Error bars denote standard deviations, which are in some cases smaller than the sizes of the symbols used in the plots. Best fit lines were calculated by fitting log_2_ transformed data. All biofilms were grown with 100 µM Nspd. All data have been corrected using the model described in the text (See Methods: Spatial correction model).

**S5 Fig. Validation of temporal and spatial QS gene expression patterns in *V. cholerae* biofilms.** (A, B, C, D) The non-corrected data used to generate the spatially corrected fluorescence signal values shown in Fig 3C, D, E, and S5E, respectively. (E) Average per-cell-volume fluorescence signal versus distance to the biofilm center, as in S3A Fig, for the data shown in Fig 3E.

**S6 Fig. Spatial QS gene expression patterns across biofilm sizes.** (A) Heatmaps showing the relative *qrr*4 (*mNG*), *hapR*, and *vpsL* per-cell-volume fluorescence signals as in Fig 2B of the main text for additional biofilm sizes. *n* = 9-10 replicate biofilms. (B) Average per-cell-volume fluorescence signal versus distance to the biofilm center is shown, as in S3A Fig, for the data represented in (A). Light symbol coloring for +AI-2 samples represents the smaller biofilms (left in (A)) and dark symbol coloring for +AI-2 samples represents larger biofilms (right in (A)). All data have been corrected using the model described in the text (See Methods: Spatial correction model).

The data for expression of *vpsL* in the AI-2 sensor strain differs markedly from that of the two upstream QS regulators in *V. cholerae* biofilms. *vpsL* exhibits changes in both its temporal and spatial expression patterns. We come back to the molecular mechanism underlying these results below, but briefly, the patterns occur because *vpsL* is regulated by both QS and by the small molecule second messenger cyclic diguanylate (c-di-GMP) [30], levels of which vary in space and time in biofilms of the AI-2 sensor strain. A few key points pertinent to the validation of our approach are relevant. First, to confirm the accuracy of measurements of *vpsL* expression, we compared *vpsL* expression levels in planktonic cells to those in the smallest biofilms in our analyses. In nascent biofilms, cells have only recently attached to the surface and begun the transition from the planktonic to the biofilm lifestyle. Thus, the biological state of *V. cholerae* cells in immature biofilms should closely resemble that of planktonic cells. Indeed, there is good agreement between *vpsL* expression levels in planktonic and early-stage biofilm cells under both LCD and HCD conditions (Fig 3A, right panel). Additionally, there is high correlation between expression levels for all three measured genes in planktonic and early-stage biofilm cells (Fig 3A, right panel, slope of best fit line on a log-log scale is 0.99). Thus, our approach also allows comparisons of expression levels across genes probed with different fluorophores. Second, unlike *qrr*4 and *hapR*, at LCD (-AI-2) *vpsL* is expressed over 2-fold more highly at the biofilm periphery than in the biofilm core, as quantified in all analyses of spatial expression (Figs 3C-E, S5E). This pattern cannot be an artifact of our spatial correction, as the pattern can be observed in raw images (Fig 3C) and is present in the non-corrected data (S5A-D Fig). Again, we explain the biology underlying formation of this pattern below. Taken together, our measurements of *hapR*, *qrr*4, and *vpsL* expression demonstrate that our smFISH strategy for probing cell-scale gene expression in *V. cholerae* biofilms can accurately quantify gene expression, both temporally and spatially.

### *V. cholerae* cells transition from the LCD to the HCD QS state as biofilms mature

Equipped with our smFISH strategy, we measured the spatiotemporal gene-expression patterns of key biofilm regulators and structural components in wildtype (WT) *V. cholerae* that undergoes the full biofilm lifecycle, from attachment to dispersal. We focused first on QS signaling and used *qrr*4 and *hapR* as readouts of the LCD and HCD QS states of cells, respectively. To capture different biofilm developmental stages, we grew biofilms for different lengths of time, ranging from six to ten hours, before fixing the samples and conducting smFISH. Consistent with previous data [32], as biofilms mature and biovolumes increase, expression of *qrr*4 decreases and expression of *hapR* increases (Fig 4A), reflecting the transition from the QS LCD-to the HCD-state. Indeed, in the largest biofilms assayed, *hapR* levels are comparable to those measured in biofilms of the AI-2 sensor strain grown with saturating AI-2 (Fig 3A, B, +AI-2). We interpret these results to mean that over the course of WT *V. cholerae* biofilm development, endogenously produced autoinducer concentrations accumulate to the level required to fully engage the QS system.

**Fig 4.**
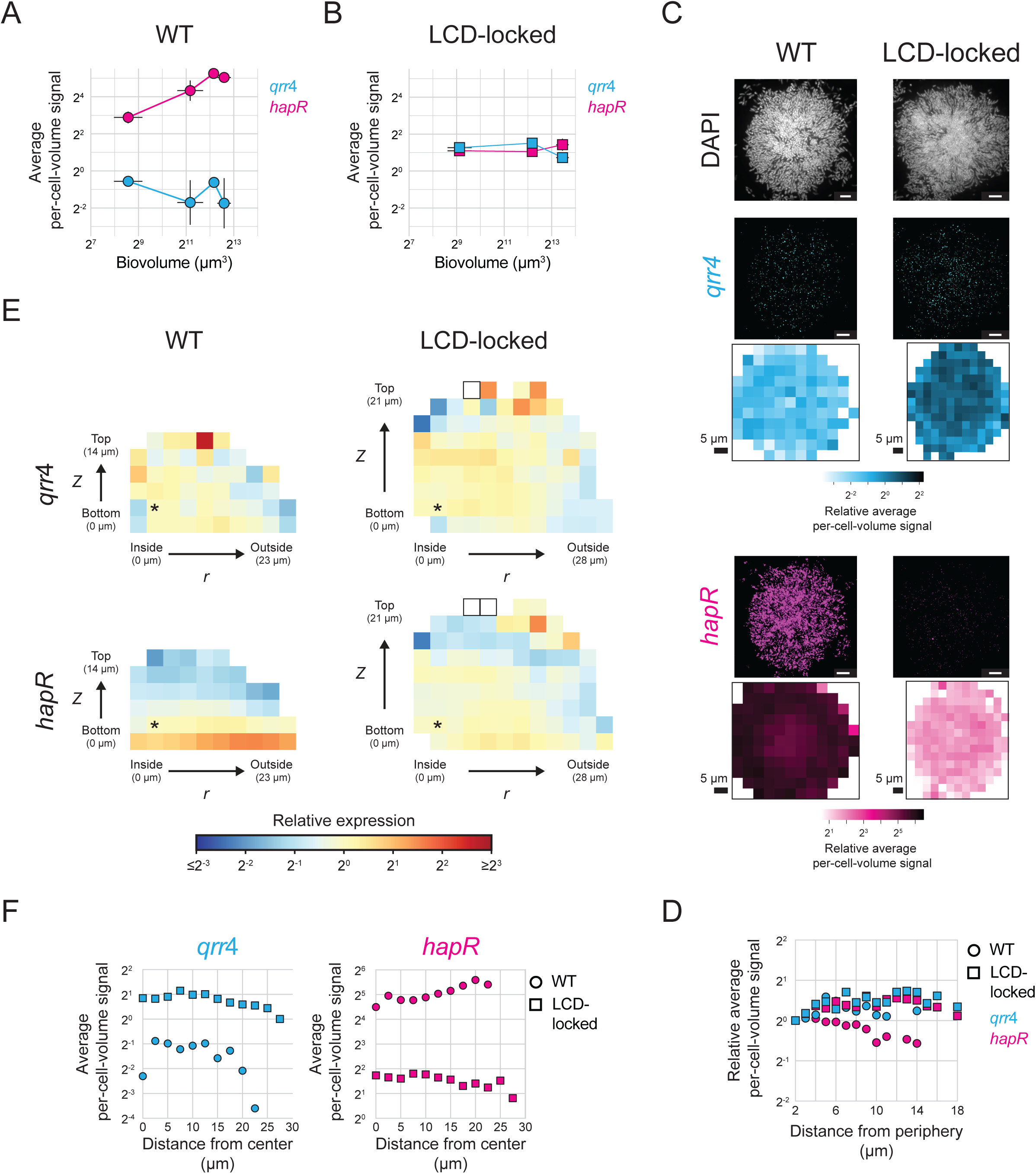
QS signaling over *V. cholerae* biofilm development. (A) Average per-cell-volume *qrr*4 (*mNG*) and *hapR* fluorescence signal across *n* = 10 replicate WT *V. cholerae* biofilms as a function of the average biofilm biovolume. (B) Average per-cell-volume *qrr*4 (*mNG*) and *hapR* fluorescence signal across *n* = 9-10 replicate *V. cholerae* biofilms formed by the LCD-locked *luxO D61E* strain as a function of the average biofilm biovolume. (C) Representative images showing DAPI, and *qrr*4 (*mNG*), and *hapR* smFISH fluorescence signals and heatmaps of per-cell-volume fluorescence signal as a function of *x*, *y* position as in Fig 3C for *n* = 9-10 replicate biofilms formed from WT (left) and LCD-locked (right) *V. cholerae* with the largest biovolumes shown in (A) and (B), respectively. Scale bars represent 5 µm. (D) Relative average per-cell-volume fluorescence signal as a function of distance from the periphery of the biofilm as in Fig 3D, for the data shown in (C). (E) Heatmaps of *qrr*4 (*mNG*) and *hapR* average per-cell-volume fluorescence signal relative to the average per-cell-volume fluorescence signal in the bins denoted with asterisks are shown as in Fig 2B for *n* = 9-10 replicate WT (left) and LCD-locked (right) *V. cholerae* biofilms with the largest biovolumes shown in (A) and (B), respectively. We note that while *z-*dependent differences in *hapR* signal are observed here, given that the *hapR* signal decreases by less than 2-fold from the surface to *z* locations far from the surface and no such pattern exists according to other spatial analyses (panels C, D, F), the differences likely stem from experiment-to-experiment variations in imaging biases that are not fully corrected by our spatial model. (F) Average per-cell-volume fluorescence signal versus distance to the biofilm center, as in S3A Fig, for the data shown in (E). Error bars denote standard deviations, which are in some cases smaller than the sizes of the symbols used in the plots. All data have been corrected using the model described in the text (See Methods: Spatial correction model).

To verify the above results, we performed control analyses in which we measured *qrr*4 and *hapR* levels over time in biofilms of *V. cholerae* carrying the constitutively active LuxO∼P phosphomimetic allele, *luxO D61E*. LuxO D61E locks cells into the LCD QS state [26]. We refer to such samples as ‘LCD-locked’. As expected, given that the LCD-locked strain cannot undergo the QS transition to HCD, expression of *qrr*4 and *hapR* remained consistent across biofilm development (Fig 4B). Furthermore, *qrr*4 expression was higher and *hapR* expression was lower in biofilms formed by the LCD-locked strain than in WT biofilms. These results confirm that the *qrr*4 and *hapR* expression changes that occur in WT *V. cholerae* biofilms are a consequence of QS. We note that *hapR* levels in the biofilms formed by the LCD-locked strain are identical to those measured in biofilms of the AI-2 sensor strain grown in the absence of AI-2 (i.e., when the AI-2 sensor strain is in the LCD QS state), however, *qrr*4 levels are lower in biofilms formed by the LCD-locked strain than in biofilms formed by AI-2 sensor strain under the no AI-2 condition (Fig 3A, B, -AI-2). We do not know the mechanism underlying this discrepancy. There exist multiple feedback loops in the QS pathway [21,45,46]. We speculate that feedback onto *qrr* expression occurring in one of these two mutant strains but not in the other could be responsible for the difference.

### QS autoinducers are spread uniformly across mature *V. cholerae* biofilms

A question that has long perplexed the biofilm field is whether QS autoinducers are present at uniform concentrations across a biofilm, or alternatively, if there exist regions of high and low autoinducer concentrations that could drive regional differences in gene-expression patterns. It is not yet possible to directly measure autoinducer concentrations in biofilms. However, the spatial accuracy that our smFISH method delivers in quantitation of gene expression in biofilms allows us to probe this longstanding question, at least in the case of *V. cholerae* biofilms. Specifically, by comparing expression levels of QS-controlled genes throughout biofilms, we can infer whether autoinducers are uniformly present or vary regionally. As expected, in biofilms containing the LCD-locked strain, spatial expression of both *qrr*4 and *hapR* differs by less than 2-fold across the biofilm (Fig 4C-F). Similar constant, location-independent expression patterns of *qrr*4 and *hapR* occur in WT biofilms, suggesting that autoinducers can freely diffuse to achieve uniform and/or saturating concentrations across the community of cells (Fig 4C-F and see text in the legend to Fig 4E regarding the *hapR* signal).

Although *qrr*4 and *hapR* do not display regional expression heterogeneity in *V. cholerae* biofilms, there is nonetheless location-independent expression variability across individual cells (S7A Fig). To investigate the source underlying this heterogeneity, we measured the relationship between *qrr*4 and *hapR* expression across cells (see Methods). Our logic is that, if location-independent non-uniformities in autoinducer concentration exist across the biofilm, the consequence would be differences in the QS states of individual cells. In that case, we would expect *qrr*4 and *hapR* expression to be negatively correlated in WT biofilms, but not in biofilms of the LCD-locked strain that is impervious to autoinducers. We do not detect any correlation between *qrr*4 and *hapR* expression in either strain, a finding consistent with intrinsic noise driving all observed heterogeneity (S7B, C Fig) [49]. We also measure negative correlations between expression variation (measured by the coefficient of variation, CV) and expression levels for both *qrr*4 and *hapR* across samples, another feature of intrinsic noise in expression (S7D Fig) [49]. This correlation is consistent across biofilms formed from the WT, LCD-locked, and AI-2 sensor strains, and between biofilm and planktonic samples, further suggesting that autoinducer fluctuations, which would be distributed and detected differently among these conditions, do not drive gene-expression heterogeneity.

**S7 Fig. Relationship between *qrr*4, *hapR*, and *vpsL* expression across biofilm cells.** (A) Cumulative distributions of per-cell-volume *qrr*4 (*mNG*) and *hapR* fluorescence signals across individual cells in WT biofilms and biofilms of the LCD-locked strain shown in Fig 4C-F. The inlay shows the region of the distribution highlighted in gray. (B) (Left) Average *qrr*4 (*mNG*) per-cell-volume fluorescence signal as a function of *hapR* per-cell-volume fluorescence signal for groups of cells in WT biofilms and biofilms of the LCD-locked strain shown in Fig 4C-F. Values calculated as described in Methods: Correlating gene expression in cell groups, using *qrr*4 as the reference gene. (Right) As on the left, with *hapR* as the reference gene. (C) As in (B) for additional biofilm sizes. Dark to light shading represents biofilms of decreasing size. (D) The relationship between the coefficient of variation (CV) for *qrr*4 (*mNG*) or for *hapR* per-cell-volume fluorescence signal across cells in individual biofilm or planktonic samples, as indicated, as a function of the average *qrr*4 (*mNG*) or *hapR* per-cell-volume signal in the sample. Empty symbols represent biofilms of the WT (circles) or of the LCD-locked strain (squares) (data from Fig 4). Slopes of the best fit lines for *qrr*4 and *hapR* for these samples are -0.42 and -0.17, respectively. Filled colored symbols represent planktonic samples of the AI-2 sensor strain grown with (circles) or without (squares) AI-2 (planktonic data from Fig 3). Slopes of the best fit lines for *qrr*4 and *hapR* for these samples are -0.43 and -0.37, respectively. Black points represent both *qrr*4 (*mNG*) and *hapR* expression in biofilms of the AI-2 sensor strain grown with or without AI-2 (biofilm data from Fig 3). Slopes of the best fit lines for *qrr*4 and *hapR* for these samples are -0.28 and -0.37, respectively. All data have been corrected using the model described in the text (See Methods: Spatial correction model).

### *vpsL* is preferentially expressed in cells located at the periphery of mature *V. cholerae* biofilms

Two important changes occur as *V. cholerae* cells transition from the planktonic lifestyle to the biofilm mode: (1) alteration in expression of hundreds of genes that enable biofilm formation and that adapt cells to the new lifestyle, and (2) organization of cells into a stereotypical yet non-uniform architecture. We hypothesized that discrete spatiotemporal patterns of expression of biofilm structural genes could arise, providing a link between gene-expression changes and specific biofilm architectural transitions. To explore this possibility, we focused first on *vpsL*, which, as mentioned, encodes a biosynthetic enzyme required for extracellular matrix polysaccharide production. *vpsL* is regulated by both QS and c-di-GMP signaling (Fig 1), allowing us to probe the contributions of both pathways to spatiotemporal gene-expression patterning. We probed biofilms formed by WT and LCD-locked *V. cholerae* to, respectively, quantify the combined contributions of QS and c-di-GMP signaling (WT) and the individual contribution of c-di-GMP signaling (LCD-locked) to establishment of the spatiotemporal *vpsL* gene-expression pattern.

Overall *vpsL* levels decrease more than 30-fold during maturation of WT biofilms (Fig 5A). We expected this temporal pattern of expression because HapR is a repressor of *vpsL*, and *hapR* is more highly expressed in mature HCD WT *V. cholerae* biofilms than in mature biofilms of the LCD-locked strain (Fig 4A, B). Temporal reductions in *vpsL* also occur in biofilms formed from the LCD-locked strain, although to a lesser extent than in WT biofilms. This temporal pattern is also observed in the AI-2 sensor strain (Fig 3B). Both the LCD-locked strain and the AI-2 sensor strain are QS inert, thus, residual changes in *vpsL* expression that occur in such biofilms cannot be a consequence of QS, but rather, must stem from changes in c-di-GMP signaling. We conclude that contributions from both QS and c-di-GMP signaling drive the temporal patterning of *vpsL* expression in WT biofilms. The reduction in *vpsL* expression that occurs as biofilms mature is likely important for terminating VPS production to allow cells to disperse from mature biofilms.

**Fig 5.**
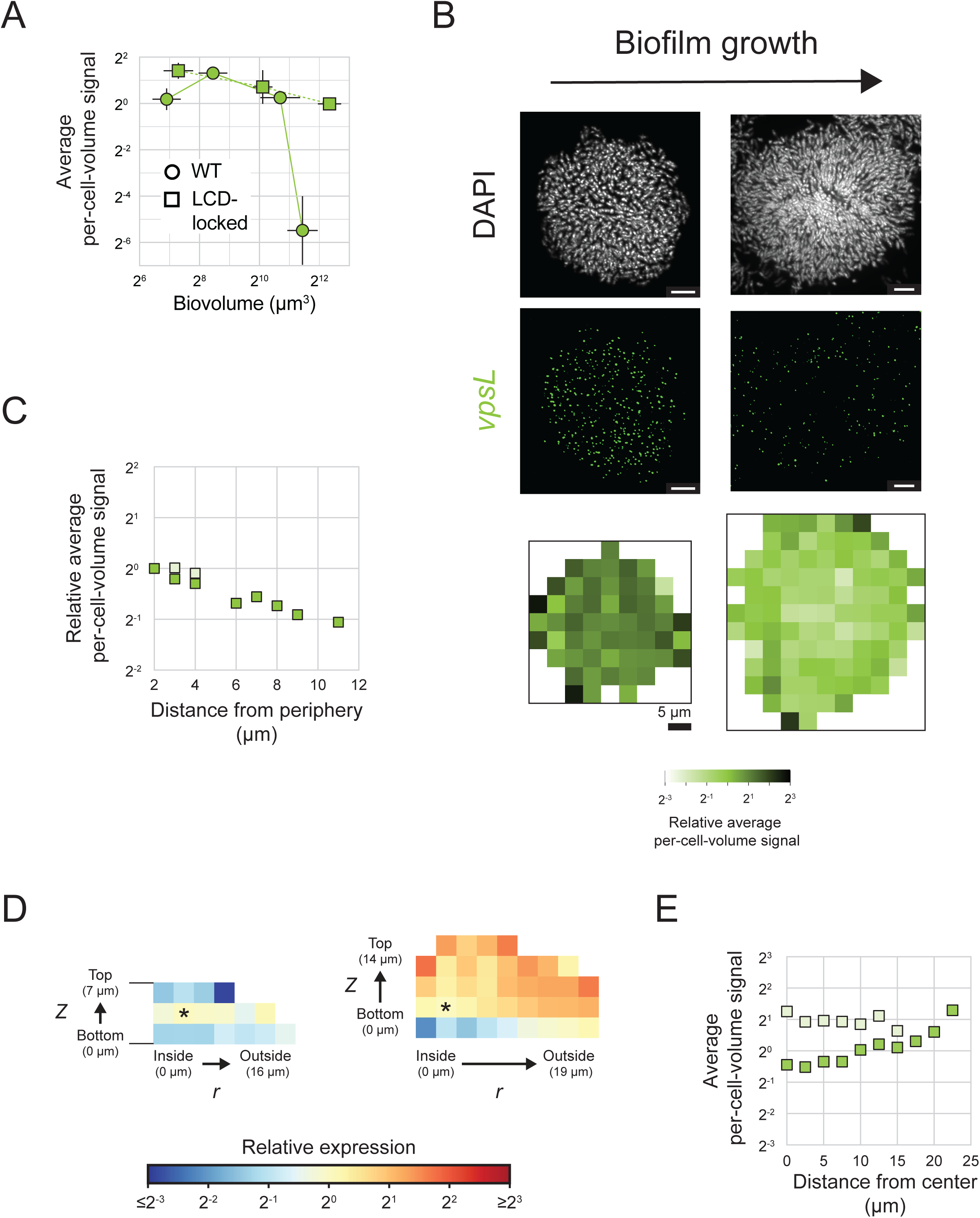
*vpsL* expression over *V. cholerae* biofilm development. (A) Average per-cell-volume *vpsL* fluorescence signal across *n* = 10 replicate WT biofilms or biofilms of the LCD-locked strain as a function of the average biofilm biovolume. Error bars represent standard deviations, which are in some cases are smaller than the sizes of the symbols used in the plots. (B) Representative images showing DAPI and *vpsL* smFISH fluorescence signals and heatmaps of per-cell-volume fluorescence signal as a function of *x*, *y* position as in Fig 3C for *n* = 10 replicate biofilms formed from LCD-locked *V. cholerae* with the two largest biovolumes shown in (A). Scale bars represent 5 µm. (C) Relative average per-cell-volume fluorescence signal as a function of distance from the periphery of the biofilm as in Fig 3D, for the data shown in (B). Light symbol coloring represents the smaller biofilms (left in (B)) and dark symbol coloring represents the larger biofilms (right in (B)). (D) Heatmaps of *vpsL* average per-cell-volume fluorescence signal relative to the average per-cell-volume fluorescence signal in the bins denoted with asterisks are shown as in Fig 2B for *n* = 10 replicate biofilms with the two largest biovolumes shown (A). (E) Average per-cell-volume fluorescence signal versus distance to the biofilm center, as in S3A Fig, for the data shown in (D). Light symbol coloring represents the smaller biofilms and dark symbol coloring represents the larger biofilms. All data have been corrected using the model described in the text (See Methods: Spatial correction model).

Our analyses show that *vpsL* expression declines to approximately 1 mRNA per 50 cells in mature WT *V. cholerae* biofilms (Fig 5A). Consequently, while we were able to monitor overall changes in *vpsL* levels by averaging over many biofilms and thousands of cells, we are unable to quantify spatial differences in *vpsL* expression in WT *V. cholerae* biofilms because, in local biofilm cell subpopulations that can contain fewer than 50 cells, we often detect zero *vpsL* mRNA transcripts. Fortuitously, mature biofilms of the LCD-locked strain, because they express overall higher levels of *vpsL* per cell than do mature WT biofilm cells, made it possible for us to probe spatial *vpsL* expression. In early-stage biofilms, *vpsL* is uniformly expressed across the biofilm (Fig 5B-E). In mature biofilms, *vpsL* transcript levels are higher in cells at the biofilm periphery than in cells located at the biofilm core (Fig 5B-E). This pattern mirrors that observed in biofilms of the AI-2 sensor strain in the absence of AI-2 (Fig 3C-E, S5E), with the difference in the magnitude of the pattern likely attributable to differences in the size, and thus developmental state, of the biofilms assayed.

Our findings regarding QS and regulation of *vpsL* spatiotemporal expression in biofilms are as follows: QS is the major driver of temporal changes in *vpsL* expression over *V. cholerae* biofilm development, and QS is responsible for terminating *vpsL* expression at biofilm maturity. While QS-independent *vpsL* expression reductions occur over time in biofilms composed of QS-deficient strains, they are modest compared to those that occur in WT biofilms. We attribute the QS-independent temporal *vpsL* expression changes to declining c-di-GMP levels that occur concurrently with biofilm maturation, as discussed below. Regarding spatial *vpsL* regulation: QS cannot establish regional differences in *vpsL* expression due to invariant autoinducer concentrations across biofilms (Fig 4C-F). Nonetheless, *vpsL* develops a distinct spatial expression pattern with higher expression in peripheral biofilm cells than in interior cells. This pattern persists in QS-inert strains (Fig 3C-E, S5E, 5B-E), further confirming the lack of a role for QS in establishing this spatial pattern. Thus, any spatial heterogeneity in *vpsL* expression that is present in biofilms must be established independently of QS activity. C-di-GMP is the only other known regulator of *vpsL*, so we conclude that regional differences in c-di-GMP signaling across the biofilm are responsible for setting up local patterns of *vpsL* gene expression. We further validate this assertion below.

### Spatiotemporal expression patterns of genes encoding matrix structural proteins parallel those of *vpsL* and are controlled by c-di-GMP signaling in *V. cholerae* biofilms

To further investigate how spatiotemporal gene-expression patterns are established during *V. cholerae* biofilm formation, we assessed the expression patterns of additional biofilm regulators and structural genes. Specifically, having considered QS regulation above, we now primarily focus on the regulators VpsR and VpsT that link alterations in c-di-GMP levels to changes in gene expression [15]. VpsR and VpsT are transcription factors that bind c-di-GMP and are active in their c-di-GMP-bound states. Once liganded, they promote expression of their downstream target genes. Transcription of *vpsT* is activated by the VpsR-c-di-GMP complex and repressed by HapR at HCD. This regulatory arrangement links *vpsT* levels to both c-di-GMP and to QS (Fig A) [30].

We measured expression of *vpsR* and *vpsT* and genes encoding the three matrix proteins RbmA, RbmC, and Bap1 that are downstream targets of VpsR and/or VpsT (Fig 1). Our smFISH technology limits us to probing three genes in a given biofilm, so we measured genes in pairs; genes assayed in the same biofilms are displayed together in the figures. We again probed both WT biofilms and biofilms formed from the LCD-locked strain to disentangle contributions from QS from those of c-di-GMP-mediated regulation. We note that the while the regulatory roles of VpsR and VpsT in biofilms are known, the timing with which and location(s) at which they act during biofilm development, to our knowledge, have not been defined.

We first established baseline expression values for *rbmA*, *rbmC*, *bap1*, *vpsR*, and *vpsT* in immature biofilms (Fig 6A). Each gene displayed a similar basal expression level in young WT biofilms and young biofilms of the LCD-locked strain. One could anticipate that the two strains would have similar expression levels given that cells in immature biofilms are in the LCD QS state. *rbmA*, *rbmC*, and *bap1* are all expressed at similar levels, which are approximately 5-fold higher than *vpsL* levels. Presumably, *vpsL,* encoding an enzyme, is not required at the quantities needed for matrix structural proteins, at least early in biofilm development. The two regulators also display striking differences in expression, with *vpsR* expressed >15-fold higher than *vpsT*.

**Fig 6.**
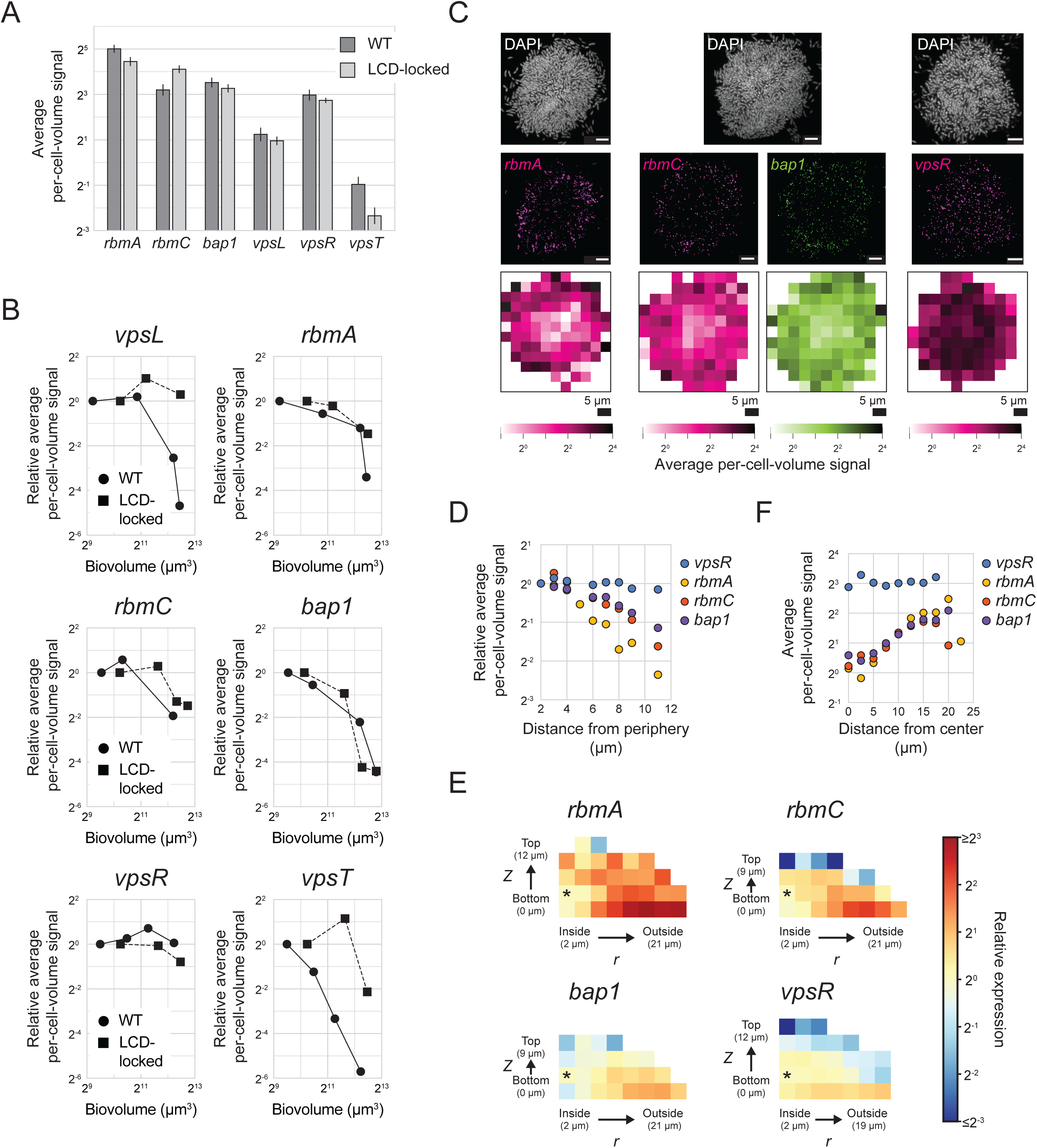
Expression of genes encoding matrix components and regulators over *V. cholerae* biofilm development. (A) Average per-cell-volume fluorescence signal of *rbmA*, *rbmC*, *bap1*, *vpsL, vpsR*, and *vpsT* across *n* = 3-7 replicate WT biofilms or biofilms formed by the LCD-locked strain, for biofilms with the smallest biovolume shown in (B). Error bars represent standard deviations. (B) Relative average per-cell-volume fluorescence signal of *vpsL*, *rbmA*, *rbmC*, *bap1*, *vpsR* and *vpsT* across *n* = 3-7 replicate WT biofilms or biofilms formed by the LCD-locked strain as a function of biovolume. Values are relative to the average per-cell-volume fluorescence signal in biofilms with the smallest biovolume for each background and gene. Pairs of plots displayed next to each other (*vpsL/rbmA*, *rbmA*/*bap1,* and *vpsR*/*vpsT*) represent pairs of genes measured in parallel in the same biofilms. (C) Representative images showing DAPI, and *rbmA*, *rbmC*, *bap1,* and *vpsR* smFISH fluorescence signals and heatmaps of per-cell-volume fluorescence signal as a function of *x*, *y* position as in Fig 3C for *n* = 5 replicate WT *V. cholerae* biofilms with the largest biovolumes shown in (B). Scale bars represent 5 µm. (D) Relative average per-cell-volume fluorescence signal as a function of distance from the periphery of the biofilm as in Fig 3D, for the data shown in (C). (E) Heatmap of *rbmA*, *rbmC*, *bap1*, and *vpsR* average per-cell-volume fluorescence signal relative to the average per-cell-volume fluorescence signal in the bin denoted with an asterisk are shown as in Fig 2B for *n* = 5 replicate WT biofilms at the largest biofilm size shown in (B). (F) Average per-cell-volume fluorescence signal versus distance to the biofilm center, as in S3A Fig, for the data shown in (E). All data have been corrected using the model described in the text (See Methods: Spatial correction model).

We hypothesized that, similar to what occurs for *vpsL* (Fig 5A), expression of the three matrix-protein encoding genes would decrease over biofilm development. Indeed, decreases do occur in each case as biofilms mature, although the timing and magnitudes of the reductions differ between the three genes (Fig 6B). In all three cases, the changes that occur in biofilms composed of the LCD-locked strain mirror those that occur in WT biofilms, indicating that their expression timing is regulated by c-di-GMP signaling and not by QS. *rbmC* and *bap1* are controlled by c-di-GMP though VpsR; they are not regulated by VpsT and they are not known to be controlled by QS. *rbmA* is controlled by c-di-GMP through VpsR and by both c-di-GMP and QS through VpsT. Consistent with regulation of *vpsT* by both QS and c-di-GMP, larger decreases in *vpsT* occur in WT biofilms than in biofilms of the LCD-locked strain (Fig 6B). Given that VpsT controls *rbmA*, it is therefore surprising that the decrease in expression of *rbmA* that occurs is equivalent in the WT and LCD-locked strains (Fig 6B). We account for this unexpected result as follows: we calculate basal expression of *vpsT* to be <1 mRNA copy per cell in both WT biofilms and in biofilms of the LCD-locked strain (Fig 6A). Likely, there is insufficient VpsT present in each biofilm cell, irrespective of the strain background or biofilm development stage, for it to act as a potent regulator of *rbmA*. Indeed, the role of VpsT in control of *V. cholerae* biofilm development has been primarily studied using a rugose variant that possess higher basal c-di-GMP levels, and consequently higher *vpsT* levels than WT and LCD-locked *V. cholerae* strains [15,50,51]. Thus, our examination of WT *V. cholerae* that possesses native c-di-GMP levels suggests that the role of VpsT in connecting c-di-GMP and QS changes to *rbmA* expression is largely inconsequential. For this reason, below we focus exclusively on the role of VpsR in connecting changes in c-di-GMP levels to changes in target gene expression in biofilms.

The reductions in *rbmA*, *rbmC*, and *bap1* that occur over biofilm development could be a consequence of decreases in c-di-GMP concentrations, decreases in *vpsR* expression, or both. *vpsR* expression remains constant over the course of biofilm formation in biofilms formed by both the WT and the LCD-locked strain (Fig 6B). These results suggest that, over the time of *V. cholerae* biofilm development, decreases in c-di-GMP levels occur that lower VpsR activity. Loss of active VpsR, in turn, results in reduced expression of the three downstream biofilm matrix genes.

One prediction stemming from our assertion that differences in c-di-GMP levels across biofilms drive the regionally-distinct pattern of *vpsL* expression is that *rbmA*, *rbmC*, and *bap1*, which we find are regulated by c-di-GMP and not QS, should behave similarly. To test this hypothesis, we measured the spatial expression patterns of the three matrix-protein encoding genes. Fortuitously, *rbmA*, *rbmC*, and *bap1* are more highly expressed than *vpsL,* enabling us to probe their spatial expression patterns in WT *V. cholerae* biofilms, which as mentioned, we could not do for *vpsL*. A clear pattern, with higher expression in peripheral cells than in cells at the core, exists for all three matrix genes in WT biofilms (Fig 6C-F). The magnitude of the spatial variability is distinct for each gene, presumably a consequence of differences in the strength of c-di-GMP-mediated regulation to which each gene is subject. The same spatial pattern of expression occurs in biofilms formed by the LCD-locked strain (S8A-E Fig), consistent with exclusive c-di-GMP regulation and no QS input into control of these three genes in either time or space in biofilms.

**S8 Fig. Spatial patterns of genes encoding matrix components in biofilms of the LCD-locked *V. cholerae* strain.** (A) Representative images showing DAPI, and *rbmA*, *rbmC*, *bap1,* and *vpsR* smFISH fluorescence signals and heatmaps of per-cell-volume fluorescence signal as a function of *x*, *y* position as in Fig 3C for *n* = 5-6 replicate *V. cholerae* biofilms formed from the LCD-locked strain with the largest biovolumes shown in Fig 6B. Scale bars represent 5 µm. (B) Relative average per-cell-volume fluorescence signal as a function of distance from the periphery of the biofilm as in Fig 3D, for the data shown in (A). (C) Representative images of *vpsR* smFISH fluorescence signal and merged DAPI and *vpsR* signal as in (A). Regions containing few cells are outlined. (D) Average per-cell-volume fluorescence signal versus distance to the biofilm center, as in S3A Fig, for the data shown in (E). (E) Heatmaps of *rbmA*, *rbmC*, *bap1*, and *vpsR* average per-cell-volume fluorescence signal relative to the average per-cell-volume fluorescence signal in the bin denoted with an asterisk are shown as in Fig 2B for *n* = 5-6 replicate biofilms formed from the LCD-locked strain with the largest biovolume shown in Fig 6B. *vpsR* appears to be preferentially expressed in cells residing at the periphery. However, the core regions of these particular biofilms contained fewer cells (see S8C) than all other biofilms analyzed in this work and consequently, there was low *vpsR* smFISH fluorescence signal output from the core. This feature biased the quantitation making it seem as if spatial pattern exists when, most likely, it does not. (F) Relationship between average per-cell-volume smFISH fluorescence signal between pairs of genes measured in parallel across groups of cells in WT biofilms and biofilms formed by the LCD-locked strain in Figs 6F and S8E, calculated as described in Methods: Correlating gene expression in cell groups. The gene on the *x*-axis represents the reference gene. (G) As in(F) for additional biofilm sizes. Dark to light shading represents biofilms of decreasing sizes. All data have been corrected using the model described in the text (See Methods: Spatial correction model).

In addition to unvarying temporal *vpsR* expression (Fig 6B), uniform *vpsR* expression also occurs across space in WT biofilms (Fig 6C-F), demonstrating that the regional patterns of expression that exist for the three VpsR target genes are not driven by regional differences in *vpsR* expression. Together, the results support our model that spatial differences in c-di-GMP levels exist across biofilm cells and, by modulating VpsR activity, establish specific spatial gene-expression patterns of target matrix genes. Further supporting the claim that differences in c-di-GMP concentration exist across cells, a positive correlation exists between expression of VpsR target genes, namely between *rbmA* and *vpsL* and between *rbmC* and *bap1,* in biofilms formed by both the WT and the LCD-locked strain (S8F, G Fig). These relationships are unlikely to result from extrinsic noise driving global cell-to-cell differences in gene expression, as no such correlation exists for other pairs of genes under study (S7B, C Fig). Rather, we suggest that the correlation stems from cell-to-cell differences in c-di-GMP concentrations driving cell-to-cell differences in VpsR activity and, in turn, expression of target genes.

### Perturbing c-di-GMP signaling disrupts the spatial patterning of *vpsL* expression in *V. cholerae* biofilms

To bolster our hypothesis that variations in c-di-GMP levels across *V. cholerae* biofilms are responsible for patterning matrix gene expression, we perturbed the endogenous c-di-GMP levels and investigated whether that altered matrix gene-expression patterns. We modulated c-di-GMP levels in biofilms by two mechanisms. First, we grew *V. cholerae* biofilms in the presence of 100 µM Nspd, which, as discussed in the Introduction, increases c-di-GMP by modulating the activity of MbaA (Fig 1). Second, we exploited a mutant *vpvC* gene, denoted *vpvC*^W240R^ and often referred to as the rugose mutation, that confers constitutive VpvC DGC activity [52]. Both Nspd addition and the *vpvC*^W240R^ mutation have been shown previously to increase cytoplasmic c-di-GMP levels, and correspondingly, matrix gene expression, with the *vpvC*^W240R^ mutation being the more potent activator of c-di-GMP production [19]. Consistent with these earlier findings, we show that *vpsL* expression is highest in biofilms formed by the LCD-locked strain containing the *vpvC*^W240R^ mutation, displays intermediate expression in biofilms of the LCD-locked strain to which Nspd has been administered, and is lowest in biofilms of the LCD-locked strain with unperturbed, endogenous c-di-GMP levels (Fig 7A). The spatial pattern in which cells at the biofilm periphery display higher *vpsL* expression than cells at the core remains present in biofilms of the LCD-locked strain following treatment with Nspd (Fig 7B-E). However, further increasing c-di-GMP levels through introduction of the *vpvC*^W240R^ mutation eliminates the *vpsL* spatial pattern (Fig 7B-E). These results suggest the following model: Globally increasing c-di-GMP levels overrides the naturally occurring differences in c-di-GMP concentrations that exist in *V. cholerae* cells in biofilms. Consequently, the c-di-GMP-controlled spatial pattern of *vpsL* gene expression is lost.

**Fig 7.**
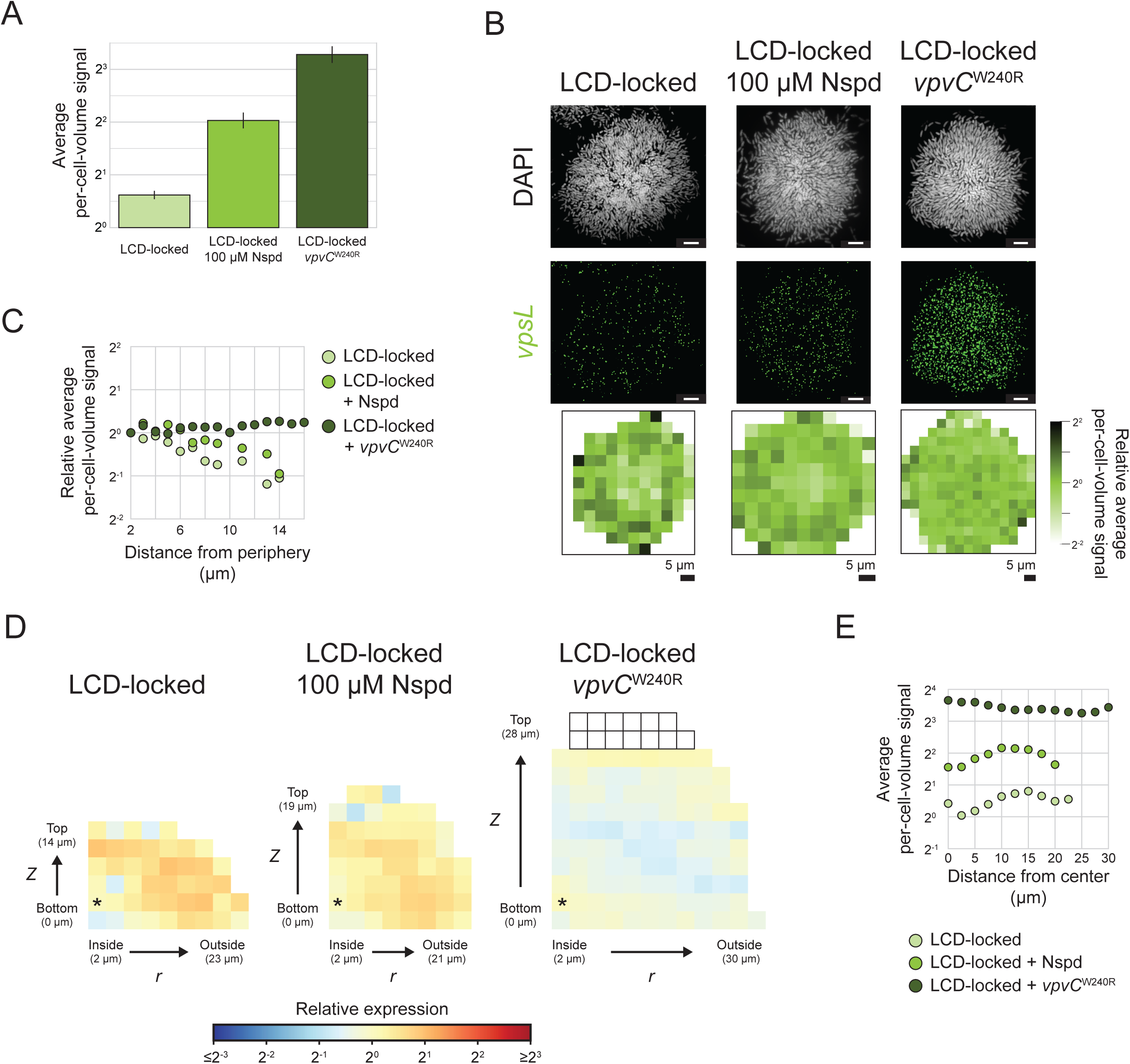
Effect of perturbation of c-di-GMP signaling on the *vpsL* spatial gene expression pattern. (A) Average per-cell-volume fluorescence signal of *vpsL* across *n* = 5 replicate biofilms formed by the LCD-locked strain, LCD-locked strain grown with 100 µM Nspd, or LCD-locked, *vpvC*^W240R^ strain. Error bars represent standard deviations. (B) Representative images showing DAPI and *vpsL* smFISH fluorescence signals, and heatmaps of per-cell-volume fluorescence signal as a function of *x*, *y* position as in Fig 3C for *n* = 5 replicate biofilms formed by the LCD-locked strain, LCD-locked strain grown with 100 µM Nspd, and LCD-locked, *vpvC*^W240R^ strain. Scale bars represent 5 µm. (C) Relative average per-cell-volume *vpsL* smFISH fluorescence signal as a function of distance from the periphery of the biofilm as in Fig 3D, for the data shown in (B). (D) Heatmaps of *vpsL* average per-cell-volume fluorescence signal relative to the average per-cell-volume fluorescence signal in the bins denoted with asterisks are shown as in Fig. 2B for *n* = 5 replicate biofilms formed by the LCD-locked strain (left), LCD-locked strain grown with 100 µM Nspd (center), or LCD-locked, *vpvC*^W240R^ strain (right). (E) Average per-cell-volume *vpsL* smFISH fluorescence signal versus distance to the biofilm center, as in S3A Fig, for the data shown in (C). All data have been corrected using the model described in the text (See Methods: Spatial correction model).

Differences in spatial *vpsL* expression patterns in biofilms of the AI-2 sensor strain grown in the presence of Nspd and in the absence or presence of AI-2 confirm the above mechanistic interpretation for how the *vpsL* biofilm spatial pattern is established. Regarding mechanism, HapR activates transcription of *nspS*-*mbaA* encoding the Nspd receptor-effector pair. Consequently, at HCD (i.e., +AI-2), when HapR is abundant, Nspd potency increases due to increased NspS-MbaA-mediated detection. Under this condition, MbaA DGC activity increases to a higher level than when Nspd is supplied in the absence of AI-2 [48]. Consistent with this mechanism, in biofilms of the AI-2 sensor strain grown with Nspd but without AI-2, *vpsL* is preferentially expressed in peripheral biofilm cells. By contrast, cells of the AI-2 sensor strain in biofilms grown with Nspd and AI-2 display uniform *vpsL* expression across space (Fig 3C-E, S5E Fig). Again, these results show that globally increasing c-di-GMP in *V. cholerae* biofilms abolishes the c-di-GMP-directed *vpsL* spatial gene-expression pattern. Moreover, the mechanism by which the c-di-GMP levels are altered is not relevant.

## Discussion

In this study, we present a biofilm-specific smFISH technology that enables accurate quantitation of spatiotemporal gene-expression patterns at cell-scale resolution and across biofilm development. This technology satisfies an urgent need in the biofilm field by providing the means to probe both the mechanistic underpinnings that drive the formation of global and regional biofilm architectures and the determinants that specify individual cell fates in biofilms. Our quantitative approach can, in principle, be applied to any gene of interest, including genes that have only low levels of expression. Our approach does not provide the throughput of other smFISH strategies such as seqFISH [41]; however, it benefits from high accuracy and individual cell resolution, expanding the types of research questions one can address. One drawback of smFISH is that it must be performed on fixed samples, which limits the temporal resolution obtainable in a single experiment. Despite this shortcoming, in this work, we were nonetheless able to reveal previously unknown spatial and temporal gene-expression patterns in *V. cholerae* biofilms. The smFISH signal decays substantially as a function of *z*, which imposes an upper bound on the thickness of biofilm samples that can be assayed (25-30 µm). In the context of the present study, *V. cholerae* cells disperse from biofilms before the communities reach 25 µm, so this issue was not a limitation for our analyses. Modifications to this technology will be required to probe thicker, non-dispersing samples.

We used spatiotemporal expression patterns of genes specifying *V. cholerae* biofilm regulators and matrix components as our test case for the smFISH technology. Our analyses reveal new information concerning how these elements are regulated in space and time. Importantly, much of the previous work on *V. cholerae* biofilm formation exploited conditions or mutants that prevented biofilm dispersal and, in so doing, interfered with native QS and/or c-di-GMP signaling [52]. These earlier investigations were seminal in that they identified key structural and regulatory components in *V. cholerae* biofilms [10,12,53], but by necessity, they prohibited study of endogenous signal transduction dynamics. While we did not focus on the biofilm dispersal process per se in this first work, our approach enabled us to study WT *V. cholerae* cells grown such that important signaling molecules that drive biofilm establishment and dispersal could vary, presumably as they do during biofilm growth in nature and during infection. Indeed, under our conditions, at late times, cells do disperse from biofilms. Our results suggest that as biofilms reach maturity, expression of genes specifying matrix components must cease in order for cells to become capable of dispersal. *V. cholerae* cells use two mechanisms to accomplish this transition. In the case of genes that are QS-repressed at HCD, namely the *vpsII* operon-encoded VPS biosynthetic genes, the timing and strength of repression is determined predominately by the HapR QS master regulator. As biofilm cells transition from the LCD to the HCD QS state, *hapR* expression increases, and *vpsII* expression is repressed. By contrast, for non-QS controlled matrix genes, the key player is c-di-GMP. Specifically, expression of genes encoding the three major matrix proteins, *rbmA*, *rbmC*, and *bap1*, is regulated exclusively by changes in c-di-GMP levels that control the activity of the transcriptional activator VpsR. For these genes, as biofilms mature, *vpsR* expression remains constant, but decreases in c-di-GMP levels result in decreased VpsR activity, and consequently suppression of the target genes.

The timing of gene-expression changes differs among the matrix genes. *bap1* expression is suppressed more strongly and earlier in the biofilm lifecycle than are *rbmA* and *rbmC*. Bap1 is important for cell-surface interactions, while RbmA and RbmC are needed for continued cell-cell and cell-matrix contacts [10]. One possible explanation for the timing differences is that Bap1 completes its primary function early in biofilm formation, such that at later times in biofilm development, lower concentrations of Bap1 suffice. Differences in the strength of regulation by c-di-GMP, through VpsR, between the promoters driving these genes likely contribute to these temporal distinctions. Future work is required to confirm this hypothesis and further connect gene-expression timing to distinct biofilm developmental events.

The smFISH approach allows measurements at cell-scale resolution. We find that quorum-sensing autoinducer levels are uniform across mature biofilms, indicating that autoinducers are able to freely diffuse through the biofilm matrix. By contrast, our analyses revealed that expression of genes encoding all the matrix components studied here is higher in cells residing at the biofilm periphery than in cells in the interior. Our data suggest this pattern is a consequence of differences in c-di-GMP levels across the biofilm, with higher c-di-GMP levels in peripheral cells driving higher VpsR activity and, thus, higher matrix component gene expression. Unfortunately, we are unable to directly measure differences in c-di-GMP concentrations in biofilm cells to bolster our mutant and imaging data because currently available reporters of c-di-GMP levels [54] are not sufficiently sensitive. However, in support of our model, our results show that this inside-to-outside spatial pattern does not depend on QS, as the pattern is preserved in biofilms of the QS-deficient, LCD-locked strain. QS is the only known regulatory pathway other than c-di-GMP signaling that feeds into control of matrix genes. Additionally, we can completely eliminate the matrix gene-expression spatial pattern by perturbing c-di-GMP levels through a variety of mechanisms.

Local differences in c-di-GMP levels that drive particular spatial and temporal patterns of biofilm matrix component production could be key to appropriately distributing tasks among individual cells in biofilm communities. We suggest that, as the biofilm matures, cells at the core have already become surrounded by extracellular matrix and so there is no need for those cells to continue to produce matrix to remain members of the biofilm community. By contrast, cells at the outside edge of the biofilm must produce and secrete matrix components to become or stay bound to the community. Further work, perhaps combining smFISH with matrix protein labeling, could connect our current findings concerning gene expression to distributions of secreted components in biofilms. Most of what is known about biofilm matrix component localization comes from field-founding studies of the rugose variant and derivatives of it [12]. As mentioned, in the rugose (VpvC^W240R^) *V. cholerae* strain, uniform expression of *vpsL* occurs across biofilms. Thus, comparisons of our data with previous findings will not yield the needed connections between gene expression and regional protein localization in biofilms, and additional studies will be necessary. Analyses focusing on times beyond those assessed in this work to monitor cells actively undergoing biofilm dispersal are also required to understand if and how spatiotemporal patterns of matrix gene expression trigger this final phase of the biofilm lifecycle.

A key finding here is that bacterial cells residing only a few microns apart experience environments sufficiently different to promote distinct modulation of their c-di-GMP levels. We currently do not know what external cue or cues are responsible for these differences. In addition to the stimuli known to modulate the activity of c-di-GMP metabolizing enzymes, for example norspermidine, sugars, and amino acids, the stimuli to which the vast majority of c-di-GMP producing/degrading enzymes respond remain unknown [18,55,56]. Thus, understanding the origin of the c-di-GMP-driven spatial patterning of matrix biofilm genes remains mysterious. Nonetheless, our smFISH approach, by enabling the spatiotemporal quantitation of genes encoding regulators, enzymes, and structural proteins provides a new and hopefully important tool for addressing questions concerning gene-expression patterns in three-dimensional living, growing bacterial communities.

## Data availability

All data, code, and raw images have been uploaded as main or supplemental files in this submission, or will be made available on Zenodo prior to publication. Additional information is available from the lead contact upon request.

## Acknowledgements

We thank members of the Bassler group for thoughtful discussion.

## Author contributions

G.E.J performed experiments; G.E.J, N.S.W, and B.L.B designed experiments and analyzed data; G.E.J and C.F wrote custom scripts for model development and image analysis; G.E.J. and B.L.B wrote the original draft; G.E.J, C.F, N.S.W, and B.L.B reviewed and edited subsequent manuscript versions. B.L.B provided oversight, resources, and funding.

## Methods

### Bacterial strains and growth conditions

The parent *V. cholerae* strain used in this study is O1 El Tor biotype C6706str2 Δ*vpsS* Δ*cqsR* (BB_Vc416). All strains used in this study are listed S1 Table. *V. cholerae* biofilms were grown as follows: Cultures were seeded from single colonies into 1 mL LB and grown at 37 °C for 3 h with shaking, to an approximate OD_600_ = 0.1. Cultures were back diluted to an OD_600_ = 6 x 10^-6^ in M9 medium (1× M9 salts, 100 μM CaCl_2_, 2 mM MgSO_4_, 0.5% dextrose, and 0.5% casamino acids). 200 µL of cultures were dispensed onto No. 1.5 glass coverslip bottomed 96-well plates (MatTek, Ashland, MA, USA), and grown statically at 37 °C. When indicated, 0.0375%, or 0.2% arabinose, 100 µM norspermidine, and/or 10 µM AI-2 was added to LB and M9 medium. When AI-2 was provided, media also contained 0.1 mM boric acid. In every case in which data for planktonic cultures are reported, the cultures were grown in the identical medium as that used for biofilm growth and the same cultures used to seed biofilm growth were used to seed planktonic cultures. Cultures were back diluted to OD_600_ = 0.0002 in 10 mL M9 medium and grown with shaking at 37 °C.

### Single-molecule RNA FISH (smFISH) probe design and hybridization

Custom Stellaris RNA FISH probes were designed against genes of interest using the Stellaris RNA FISH Probe Designer v.4.2 (LGC, Biosearch Technologies). Probes were labeled with Quasar 670, CAL Fluor Red□590, or FAM (see S2 Table for a complete list of probe sequences and associated fluorophores). For biofilm samples, probes were hybridized following the manufacturer’s instructions for adherent cells in 96-well glass bottom plates, with minor modifications. The full protocol is available online at www.biosearchtech.com/stellarisprotocols. Briefly, following growth, medium was decanted, and adhered biofilms were fixed for 10 min at room temperature with 200 µL 1x PBS and 3.7% formaldehyde. The fixed biofilms were washed four times with 200 µL 1x PBS. To permeabilize cells, the samples were incubated overnight, or up to 24 h, at 4 °C in 200 µL 70% EtOH. Samples were incubated for 5 min at room temperature with 200 µL RNA FISH wash buffer□A (Biosearch Technologies, prepared following the manufacturer’s instructions) containing 10% formamide. Probes were hybridized with 75 µL RNA FISH hybridization buffer (Biosearch Technologies) containing 10% formamide and 1.5□µL of each probe stock (12.5□µM), followed by overnight incubation at 37□°C. Samples were washed twice for 30 min at 37 °C with 200 µL RNA FISH wash buffer□A containing 10% formamide. Cells were stained with 50□µg□mL^−1^ DAPI in wash buffer□A for 20□min at 37□°C. Following a final wash with 200□µL Stellaris RNA FISH wash buffer□B, 50□µL VectaShield mounting medium was added, and the samples were immediately imaged. Regarding planktonic cells, probe hybridization was performed as previously described [39], with minor modifications as detailed in [57].

### Confocal imaging

Imaging was performed with a Nikon Eclipse Ti2 inverted microscope equipped with a Yokogawa CSU-W1 SoRa confocal scanning unit. Samples were imaged with a CFI Apochromat TIRF ×60 oil objective lens (Nikon, 1.49 numerical aperture) with excitation wavelengths of 405 (DAPI), 488 (mNG and FAM), 561 (mScarlet and CAL Fluor Red 590), and 640□nm (Quasar 670) and with 0.5□µm *z*-steps. Images were captured through a ×2.8 SoRa magnifier.

### smFISH image analysis

Images were processed using Nikon NIS-Elements Denoise.ai software. Denoised images of planktonic cells were segmented using Fiji software [58] from maximum *z*-projections of the DAPI channel. A gaussian blur was applied to images, and cells were identified using an intensity threshold. Cells that could not be accurately resolved from neighboring cells were excluded from downstream analyses. Using the DAPI channel, denoised images of biofilm cells were segmented with BiofilmQ into cubes with side lengths of 2.32 µm [43]. The center *x, y, z* coordinates of each cell cube and the volume fraction of each cube occupied by cell mass were exported for use in downstream analyses. We interchangeably use “cell cube” and “cell” to refer to this resulting cube-based segmentation. The boundaries of individual biofilms in denoised images were manually defined using the DAPI channel. Cell cubes outside of these boundaries were excluded from downstream analysis.

smFISH data were analyzed using custom python scripts. Briefly, images were convolved with a Gaussian function to remove noise. Puncta were detected as local maxima with intensities greater than a threshold that was established based on a set of negative control images specific to each probe set. For *hapR* and *mNG* probes, negative control images were acquired using biofilms containing cells lacking *hapR* and *mNG* (BBVc_801). For *vpsL* probes, mature WT biofilms (BBVc_416), i.e., at a stage when *vpsL* is no longer expressed, were hybridized with *vpsL* to acquire negative control data. The *vpsR, vpsT, rbmA, rbmC,* and *bap1* genes could not be deleted, because without them, biofilms do not form. Likewise, *polA* and *gyrA* are essential genes and cannot be deleted. Thus, as a negative control for these probe sets, WT (BBVc_416) biofilms, i.e., biofilms lacking *mNG*, were hybridized with *mNG*-targeting probes labeled with the fluorophore matching each probe set of interest. Each punctum was fitted with a 3D Gaussian function to determine the integrated, background-subtracted punctum intensity. In instances with multiple puncta residing in close proximity, a 3D multi-Gaussian fit was performed. Puncta were assigned to cells (planktonic samples) or to cell cubes (biofilm samples) and the total fluorescence signal per cell or cube was calculated as the sum of all puncta intensities. For biofilm samples, this fluorescence signal was further normalized by the Cube_VolumeFraction, which is the fraction of the cube volume occupied by biomass, calculated by BiofilmQ.

### Spatial correction model

Spatial correction factors were calculated using data from the largest biofilms formed by the parent strain carrying Δ*vc1807*::pBad-*mNG* (BB_Vc813) imaged under each condition. Unique correction factors were derived for each of the three fluorophores, at both high and low expression levels of *mNG*, and for all positions (*r, z*). As biofilms are not perfect half spheres, the *r* position of each cell cube was calculated in a *z*-dependent manner. Specifically, a unique center position was determined for each *z* bin as the volume-weighted average *x*, *y* coordinate across all cells within that *z* bin. For *z* bins with *z* ≤ 391 pixels (15.1 µm) and *z* > 391 pixels (15.1 µm), this procedure shifted the calculated center position relative to the *x, y* center of the biofilm as a whole by an average of 4.8 µm (SD = 2.6 µm) and 13.0 µm (SD = 5.6 µm), respectively. The *r* position for each cell cube was subsequently calculated as the distance between the cell cube’s *x, y* location and the center *x, y* position corresponding to that cell cube’s *z* location. The correction factors use experimental data to describe both *z* and *r* positional biases, as detailed below. To calculate *z* bias, the average per-cell-volume smFISH fluorescence signals from cell cubes with radial position *r* = 311 pixels (12.0 µm) were calculated as a function of cell cube *z* position and normalized to the average per-cell-volume smFISH fluorescence signal of cell cubes with *z, r =* 91, 311 pixels (3.52, 12.0 µm). These data were fit with the ad-hoc Equation (1) to determine the correction factor as a function of *z*:

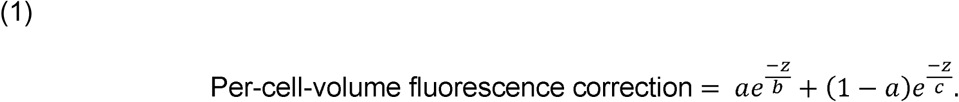

To calculate *r* bias, the average per-cell-volume fluorescence signal of cell cubes with *z* position *z* = 91 pixels (3.52 µm) were calculated as a function of cell cube *r* position. We observed that the average smFISH fluorescence signal decreased linearly as a function of *r* before plateauing at *r* = *r_i_*, with the position of *r_i_* differing for each sample. For each of the six samples, we chose an *r_i_* value by visual inspection of the data. The average smFISH fluorescence signal was normalized to the average smFISH fluorescence signal of cell cubes with *r* = *r_i_*, *z* = 91 pixels (3.52 µm) and fit to the ad-hoc Equation (2):

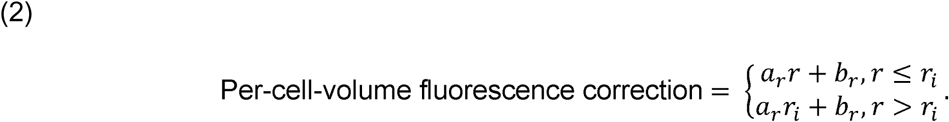

Using the specific fit parameters for each fluorophore and for high and low gene-expression levels, the correction factor for all positions (*r, z*) was calculated for each condition as the product of Equation (1) and Equation (2). These values form the basis for calculating interpolative equations used to correct all datasets, as described below.

In order to acquire appropriate correction factors for smFISH fluorescent signals from genes with any expression level, not only those matching the *mNG* high and low expression conditions, and also for biofilms of any size, we developed a generalized interpolation model as follows: for a given fluorophore, for each position (*r, z*), a linear equation *Y*(*X*) connecting points (*X*_1_, *Y*_1_) and (*X*_2_, *Y*_2_) was calculated, in which *Y*_1_ and *Y*_2_ are the correction factors derived above from the low-and high-gene-expression conditions, respectively, and *X*_1_ and *X*_2_ are the log_10_ values of the average summed smFISH fluorescence signals across the biofilms that were used to calculate the above correction factors in the low-and high-gene-expression conditions, respectively. With this model, for any given biofilm or group of replicate biofilms, unique correction factors can be determined for each position (*r, z*) based on the total smFISH fluorescent signal in the biofilm(s). Specifically, for each position (*r, z*), the log_10_ of the average summed smFISH fluorescence signals across replicate biofilms was taken as an *X* value and introduced into the corresponding (*r, z*)-dependent linear equation *Y*(*X*) generated above to calculate an (*r, z*)-specific correction factor for those biofilms. To use these calculated correction factors, for each cell cube within the biofilm, the per-cell-volume smFISH fluorescence signal, calculated as described above, was divided by the correction factor based on its position (*r, z*). The resulting calculated values were used for all subsequent analyses, unless otherwise noted. Values for all fit parameters, (*r, z*)-dependent correction factors, and resulting (*r, z*)-dependent interpolative linear equations can be found in S3 Table. The code detailing generation and application of the model, and accompanying sample plots of fit parameters, will be deposited and available prior to publication.

### Analysis of spatial gene-expression patterns at cell-scale resolution

Spatial gene-expression patterns in individual cells were assessed in several ways. First, to visualize radial patterns (as in Fig 3C), cell cubes at the bottom of the biofilm, defined as cells with *z* positions between 0 and 4.6 µm, across replicate biofilms were grouped by their *x* and *y* positions. For each cell cube, the *x, y* position was set relative to the center *x, y* position of the biofilm. The per-cell-volume signal for each group of cells was calculated and plotted as heatmaps using bins with Δ*x*, Δ*y* = 100, 100 pixels (3.9 µm, 3.9 µm). Second, the distance of each cell cube at the bottom of the biofilm to the periphery of the biofilm was calculated as the shortest distance from the cell to the biofilm perimeter, which was manually defined. Based on this distance, cells across replicate biofilms were grouped into 1 µm sized bins and the relative average per-cell-volume fluorescence signal per bin was calculated relative to cells with distance 1 µm from the biofilm periphery. Resulting data were plotted as a function of the left boundary of each bin (as in Fig 3D) for bins with greater than n = sample size x 10 cells. Third, cell cubes across replicate biofilms were grouped by their center *r* and *z* positions. Radial positions for each cell cube were calculated as described above. The average per-cell-volume fluorescence signal for each group of cells was calculated and normalized to the group of cells with center position *r, z* = 91, 91 pixels (3.5 µm, 3.5 µm). Resulting data for groups containing ≥ 25 cells were plotted as heatmaps using bins with Δ*r,* Δ *z* = 2.32, 2.32 µm (as in Fig 2). Fourth, the distance of each cell cube from the center of the biofilm was calculated from the center *r, z* position of that cell cube to the center position *r, z* = 0, 91 pixels (0, 3.5 µm). Based on this calculated distance, cells across replicate biofilms were grouped into 2.5 µm sized bins, and the average per-cell-volume fluorescence signal per bin was calculated. Resulting data were plotted as a function of the left boundary of each bin (as in S3A Fig).

### Correlating gene expression in cell groups

The relationship between gene-expression levels measured in parallel across cells was assessed as follows: For a given ‘reference gene’, all cells with zero signal corresponding to that gene were omitted from analysis. The remaining cells were ordered by expression level of the reference gene and grouped into bins of 500 cells each. The average per-cell-volume fluorescence signal was calculated for each bin for both the reference gene and a second ‘comparison gene’, and these two average signal values were plotted against one another for each bin.

### Quantitation and statistical analyses

Standard deviations were calculated using custom Python scripts from log_2_ transformed data.

*z*-scores of average per-cell-volume fluorescence signal in *r, z* bins were calculated using the following formula, 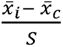 where *x̅*_i_ represents the mean per-cell-volume fluorescence signal across cells in the bin, *S* is the standard deviation of the per-cell-volume fluorescence signal across cells in the bin, and *x̅*_c_ is the mean per-cell-volume fluorescence signal in the bin with *r, z* = 91, 91 pixels (3.5 µm, 3.5 µm).

---

**S1 Table. Strains used in this study.**

**S2 Table. smFISH probe sequences used in this study.**

**S3 Table. Values of fitting and correction parameters generated by the spatial correction model.**

**S1 Data. Companion plots generated from non-corrected data.** For each plot, the corresponding main text or supplemental figure is indicated.

## References

1. Flemming H-C, Wingender J, Szewzyk U, Steinberg P, Rice SA, Kjelleberg S. Biofilms: an emergent form of bacterial life. Nat Rev Microbiol. 2016;14: 563–575. doi:10.1038/nrmicro.2016.94

2. Mah TF, O’Toole GA. Mechanisms of biofilm resistance to antimicrobial agents. Trends Microbiol. 2001;9: 34–39. doi:10.1016/s0966-842x(00)01913-2

3. Shaw T, Winston M, Rupp CJ, Klapper I, Stoodley P. Commonality of Elastic Relaxation Times in Biofilms. Phys Rev Lett. 2004;93: 098102. doi:10.1103/PhysRevLett.93.098102

4. Nadell CD, Drescher K, Wingreen NS, Bassler BL. Extracellular matrix structure governs invasion resistance in bacterial biofilms. ISME J. 2015;9: 1700–1709. doi:10.1038/ismej.2014.246

5. Flemming H-C, Wingender J. The biofilm matrix. Nat Rev Microbiol. 2010;8: 623–633. doi:10.1038/nrmicro2415

6. Guilhen C, Forestier C, Balestrino D. Biofilm dispersal: multiple elaborate strategies for dissemination of bacteria with unique properties. Molecular Microbiology. 2017;105: 188–210. doi:10.1111/mmi.13698

7. Bridges AA, Fei C, Bassler BL. Identification of signaling pathways, matrix-digestion enzymes, and motility components controlling Vibrio cholerae biofilm dispersal. Proceedings of the National Academy of Sciences. 2020;117: 32639–32647. doi:10.1073/pnas.2021166117

8. Conner JG, Teschler JK, Jones CJ, Yildiz FH. Staying alive: Vibrio cholerae’s cycle of environmental survival, transmission, and dissemination. Microbiol Spectr. 2016;4:10.1128/microbiolspec.VMBF-0015-2015

9. Gallego-Hernandez AL, DePas WH, Park JH, Teschler JK, Hartmann R, Jeckel H, et al. Upregulation of virulence genes promotes Vibrio cholerae biofilm hyperinfectivity. PNAS. 2020;117: 11010–11017. doi:10.1073/pnas.1916571117

10. Teschler JK, Zamorano-Sánchez D, Utada AS, Warner CJA, Wong GCL, Linington RG, et al. Living in the matrix: assembly and control of Vibrio cholerae biofilms. Nat Rev Microbiol. 2015;13: 255–268. doi:10.1038/nrmicro3433

11. Teschler JK, Nadell CD, Drescher K, Yildiz FH. Mechanisms Underlying Vibrio cholerae Biofilm Formation and Dispersion. Annual Review of Microbiology. 2022;76: 503–532. doi:10.1146/annurev-micro-111021-053553

12. Berk V, Fong JCN, Dempsey GT, Develioglu ON, Zhuang X, Liphardt J, et al. Molecular Architecture and Assembly Principles of Vibrio cholerae Biofilms. Science. 2012;337: 236–239. doi:10.1126/science.1222981

13. Huang X, Nero T, Weerasekera R, Matej KH, Hinbest A, Jiang Z, et al. Vibrio cholerae biofilms use modular adhesins with glycan-targeting and nonspecific surface binding domains for colonization. Nat Commun. 2023;14: 2104. doi:10.1038/s41467-023-37660-0

14. Hammer BK, Bassler BL. Quorum sensing controls biofilm formation in Vibrio cholerae. Mol Microbiol. 2003;50: 101–104. doi:10.1046/j.1365-2958.2003.03688.x

15. Zamorano-Sánchez D, Fong JCN, Kilic S, Erill I, Yildiz FH. Identification and Characterization of VpsR and VpsT Binding Sites in Vibrio cholerae. Journal of Bacteriology. 2015;197: 1221–1235. doi:10.1128/jb.02439-14

16. Srivastava D, Harris RC, Waters CM. Integration of Cyclic di-GMP and Quorum Sensing in the Control of vpsT and aphA in Vibrio cholerae Ill. J Bacteriol. 2011;193: 6331–6341. doi:10.1128/JB.05167-11

17. Conner JG, Zamorano-Sánchez D, Park JH, Sondermann H, Yildiz FH. The ins and outs of cyclic di-GMP signaling in Vibrio cholerae. Curr Opin Microbiol. 2017;36: 20–29. doi:10.1016/j.mib.2017.01.002

18. Bridges AA, Bassler BL. Inverse regulation of Vibrio cholerae biofilm dispersal by polyamine signals. Laub MT, Storz G, Michael A, Mandel M, editors. eLife. 2021;10: e65487. doi:10.7554/eLife.65487

19. Bridges AA, Prentice JA, Fei C, Wingreen NS, Bassler BL. Quantitative input–output dynamics of a c-di-GMP signal transduction cascade in Vibrio cholerae. PLOS Biology. 2022;20: e3001585. doi:10.1371/journal.pbio.3001585

20. Mukherjee S, Bassler BL. Bacterial quorum sensing in complex and dynamically changing environments. Nat Rev Microbiol. 2019;17: 371–382. doi:10.1038/s41579-019-0186-5

21. Bridges AA, Prentice JA, Wingreen NS, Bassler BL. Signal Transduction Network Principles Underlying Bacterial Collective Behaviors. Annual Review of Microbiology. 2022;76: 235–257. doi:10.1146/annurev-micro-042922-122020

22. Higgins DA, Pomianek ME, Kraml CM, Taylor RK, Semmelhack MF, Bassler BL. The major Vibrio cholerae autoinducer and its role in virulence factor production. Nature. 2007;450: 883–886. doi:10.1038/nature06284

23. Wei Y, Perez LJ, Ng W-L, Semmelhack MF, Bassler BL. Mechanism of Vibrio cholerae Autoinducer-1 Biosynthesis. ACS Chem Biol. 2011;6: 356–365. doi:10.1021/cb1003652

24. Zhao G, Wan W, Mansouri S, Alfaro JF, Bassler BL, Cornell KA, et al. Chemical synthesis of *S*-ribosyl-l-homocysteine and activity assay as a LuxS substrate. Bioorganic & Medicinal Chemistry Letters. 2003;13: 3897–3900. doi:10.1016/j.bmcl.2003.09.015

25. Neiditch MB, Federle MJ, Miller ST, Bassler BL, Hughson FM. Regulation of LuxPQ receptor activity by the quorum-sensing signal autoinducer-2. Mol Cell. 2005;18: 507–518. doi:10.1016/j.molcel.2005.04.020

26. Freeman JA, Bassler BL. A genetic analysis of the function of LuxO, a two-component response regulator involved in quorum sensing in Vibrio harveyi. Molecular Microbiology. 1999;31: 665–677. doi:10.1046/j.1365-2958.1999.01208.x

27. Freeman JA, Bassler BL. Sequence and Function of LuxU: a Two-Component Phosphorelay Protein That Regulates Quorum Sensing inVibrio harveyi. Journal of Bacteriology. 1999;181: 899–906. doi:10.1128/jb.181.3.899-906.1999

28. Lenz DH, Mok KC, Lilley BN, Kulkarni RV, Wingreen NS, Bassler BL. The Small RNA Chaperone Hfq and Multiple Small RNAs Control Quorum Sensing in Vibrio harveyi and Vibrio cholerae. Cell. 2004;118: 69–82. doi:10.1016/j.cell.2004.06.009

29. Rutherford ST, van Kessel JC, Shao Y, Bassler BL. AphA and LuxR/HapR reciprocally control quorum sensing in vibrios. Genes Dev. 2011;25: 397–408. doi:10.1101/gad.2015011

30. Waters CM, Lu W, Rabinowitz JD, Bassler BL. Quorum sensing controls biofilm formation in Vibrio cholerae through modulation of cyclic di-GMP levels and repression of vpsT. J Bacteriol. 2008;190: 2527–2536. doi:10.1128/JB.01756-07

31. Yildiz FH, Liu XS, Heydorn A, Schoolnik GK. Molecular analysis of rugosity in a Vibrio cholerae O1 El Tor phase variant. Molecular Microbiology. 2004;53: 497–515. doi:10.1111/j.1365-2958.2004.04154.x

32. Bridges AA, Bassler BL. The intragenus and interspecies quorum-sensing autoinducers exert distinct control over Vibrio cholerae biofilm formation and dispersal. PLoS Biol. 2019;17: e3000429. doi:10.1371/journal.pbio.3000429

33. Drescher K, Dunkel J, Nadell CD, van Teeffelen S, Grnja I, Wingreen NS, et al. Architectural transitions in Vibrio cholerae biofilms at single-cell resolution. Proceedings of the National Academy of Sciences. 2016;113: E2066–E2072. doi:10.1073/pnas.1601702113

34. Yan J, Sharo AG, Stone HA, Wingreen NS, Bassler BL. Vibrio cholerae biofilm growth program and architecture revealed by single-cell live imaging. PNAS. 2016;113: E5337– E5343. doi:10.1073/pnas.1611494113

35. Qin B, Fei C, Bridges AA, Mashruwala AA, Stone HA, Wingreen NS, et al. Cell position fates and collective fountain flow in bacterial biofilms revealed by light-sheet microscopy. Science. 2020;369: 71–77. doi:10.1126/science.abb8501

36. Heim R, Prasher DC, Tsien RY. Wavelength mutations and posttranslational autoxidation of green fluorescent protein. Proc Natl Acad Sci U S A. 1994;91: 12501–12504. doi:10.1073/pnas.91.26.12501

37. Singh PK, Bartalomej S, Hartmann R, Jeckel H, Vidakovic L, Nadell CD, et al. Vibrio cholerae Combines Individual and Collective Sensing to Trigger Biofilm Dispersal. Current Biology. 2017;27: 3359–3366.e7. doi:10.1016/j.cub.2017.09.041

38. Manner C, Dias Teixeira R, Saha D, Kaczmarczyk A, Zemp R, Wyss F, et al. A genetic switch controls Pseudomonas aeruginosa surface colonization. Nat Microbiol. 2023;8: 1520–1533. doi:10.1038/s41564-023-01403-0

39. Skinner SO, Sepúlveda LA, Xu H, Golding I. Measuring mRNA copy number in individual Escherichia coli cells using single-molecule fluorescent in situ hybridization. Nat Protoc. 2013;8: 1100–1113. doi:10.1038/nprot.2013.066

40. So L, Ghosh A, Zong C, Sepúlveda LA, Segev R, Golding I. GENERAL PROPERTIES OF THE TRANSCRIPTIONAL TIME-SERIES IN ESCHERICHIA COLI. Nat Genet. 2011;43: 554–560. doi:10.1038/ng.821

41. Dar D, Dar N, Cai L, Newman DK. Spatial transcriptomics of planktonic and sessile bacterial populations at single-cell resolution. Science. 2021;373: eabi4882. doi:10.1126/science.abi4882

42. Espinosa E, Daniel S, Hernández SB, Goudin A, Cava F, Barre F-X, et al. l-Arabinose Induces the Formation of Viable Nonproliferating Spheroplasts in Vibrio cholerae. Appl Environ Microbiol. 2021;87: e02305–20. doi:10.1128/AEM.02305-20

43. Hartmann R, Jeckel H, Jelli E, Singh PK, Vaidya S, Bayer M, et al. Quantitative image analysis of microbial communities with BiofilmQ. Nat Microbiol. 2021;6: 151–156. doi:10.1038/s41564-020-00817-4

44. Papenfort K, Förstner KU, Cong J-P, Sharma CM, Bassler BL. Differential RNA-seq of Vibrio cholerae identifies the VqmR small RNA as a regulator of biofilm formation. PNAS. 2015;112: E766–E775. doi:10.1073/pnas.1500203112

45. Svenningsen SL, Waters CM, Bassler BL. A negative feedback loop involving small RNAs accelerates Vibrio cholerae’s transition out of quorum-sensing mode. Genes Dev. 2008;22: 226–238. doi:10.1101/gad.1629908

46. Svenningsen SL, Tu KC, Bassler BL. Gene dosage compensation calibrates four regulatory RNAs to control Vibrio cholerae quorum sensing. The EMBO Journal. 2009;28: 429–439. doi:10.1038/emboj.2008.300

47. Neiditch MB, Federle MJ, Pompeani AJ, Kelly RC, Swem DL, Jeffrey PD, et al. Ligand-induced asymmetry in histidine sensor kinase complex regulates quorum sensing. Cell. 2006;126: 1095–1108. doi:10.1016/j.cell.2006.07.032

48. Prentice JA, Bridges AA, Bassler BL. Synergy between c-di-GMP and Quorum-Sensing Signaling in Vibrio cholerae Biofilm Morphogenesis. Journal of Bacteriology. 2022;204: e00249–22. doi:10.1128/jb.00249-22

49. Raj A, van Oudenaarden A. Stochastic gene expression and its consequences. Cell. 2008;135: 216–226. doi:10.1016/j.cell.2008.09.050

50. Casper-Lindley C, Yildiz FH. VpsT is a transcriptional regulator required for expression of vps biosynthesis genes and the development of rugose colonial morphology in Vibrio cholerae O1 El Tor. J Bacteriol. 2004;186: 1574–1578. doi:10.1128/JB.186.5.1574-1578.2004

51. Beyhan S, Bilecen K, Salama SR, Casper-Lindley C, Yildiz FH. Regulation of Rugosity and Biofilm Formation in Vibrio cholerae: Comparison of VpsT and VpsR Regulons and Epistasis Analysis of vpsT, vpsR, and hapR. Journal of Bacteriology. 2007;189: 388–402. doi:10.1128/jb.00981-06

52. Beyhan S, Yildiz FH. Smooth to rugose phase variation in Vibrio cholerae can be mediated by a single nucleotide change that targets c-di-GMP signalling pathway. Molecular Microbiology. 2007;63: 995–1007. doi:10.1111/j.1365-2958.2006.05568.x

53. Fong JCN, Syed KA, Klose KE, Yildiz FH. Role of Vibrio polysaccharide (vps) genes in VPS production, biofilm formation and Vibrio cholerae pathogenesis. Microbiology (Reading). 2010;156: 2757–2769. doi:10.1099/mic.0.040196-0

54. Zhou H, Zheng C, Su J, Chen B, Fu Y, Xie Y, et al. Characterization of a natural triple-tandem c-di-GMP riboswitch and application of the riboswitch-based dual-fluorescence reporter. Sci Rep. 2016;6: 20871. doi:10.1038/srep20871

55. Heo K, Park Y-H, Lee K-A, Kim J, Ham H-I, Kim B-G, et al. Sugar-mediated regulation of a c-di-GMP phosphodiesterase in Vibrio cholerae. Nat Commun. 2019;10: 5358. doi:10.1038/s41467-019-13353-5

56. Xu M, Wang Y-Z, Yang X-A, Jiang T, Xie W. Structural studies of the periplasmic portion of the diguanylate cyclase CdgH from Vibrio cholerae. Sci Rep. 2017;7: 1861. doi:10.1038/s41598-017-01989-6

57. Silpe JE, Duddy OP, Johnson GE, Beggs GA, Hussain FA, Forsberg KJ, et al. Small protein modules dictate prophage fates during polylysogeny. Nature. 2023;620: 625–633. doi:10.1038/s41586-023-06376-y

58. Schindelin J, Arganda-Carreras I, Frise E, Kaynig V, Longair M, Pietzsch T, et al. Fiji: an open-source platform for biological-image analysis. Nat Methods. 2012;9: 676–682. doi:10.1038/nmeth.2019

