## Supplementary figures and images for "Cell-scale gene-expression measurements in *Vibrio cholerae* biofilms reveal spatiotemporal patterns underlying development"

### Supplemental Figures

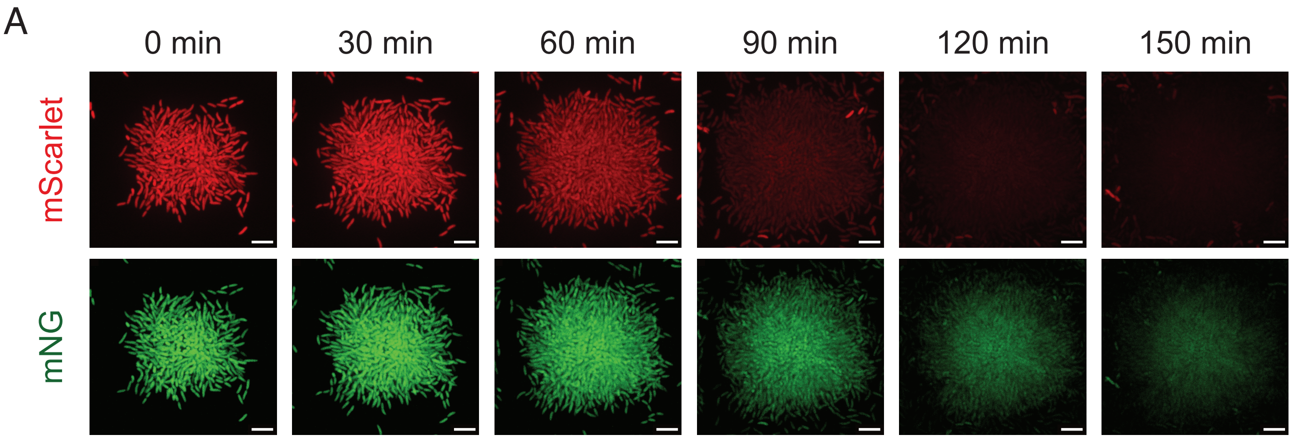

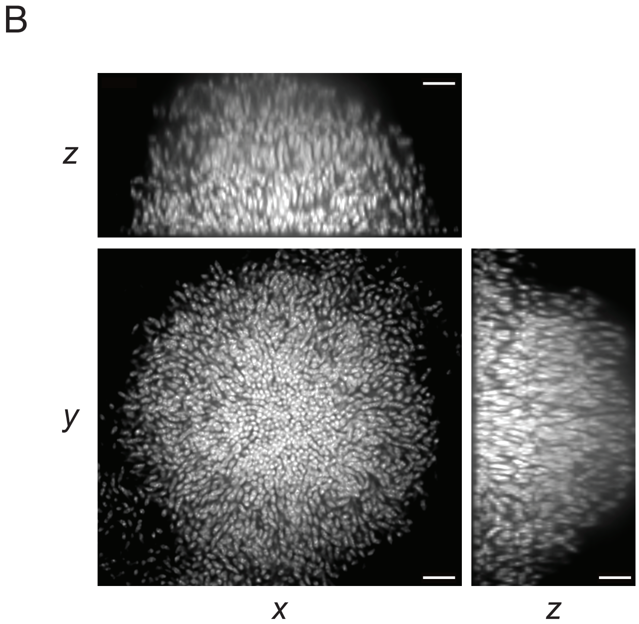

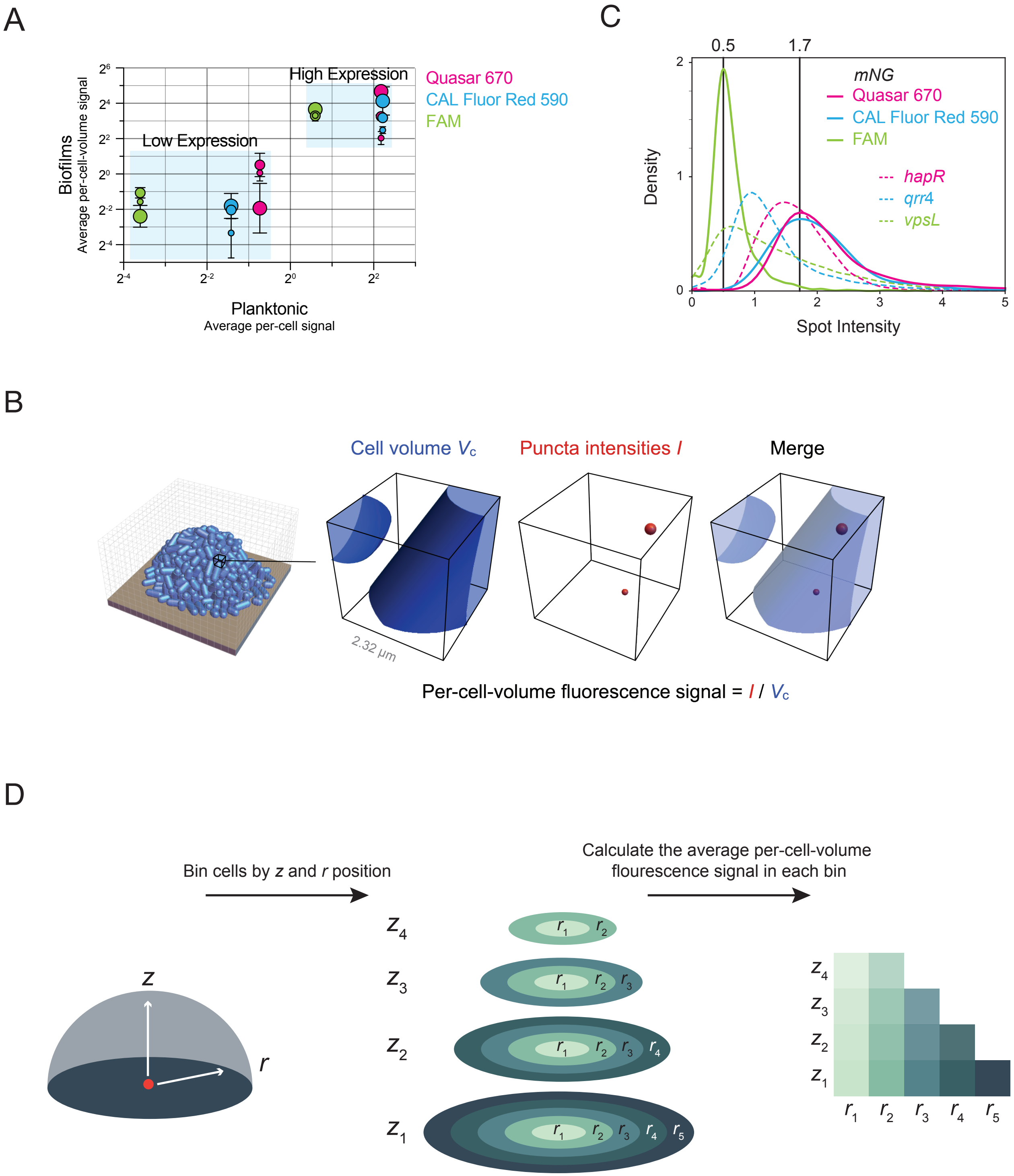

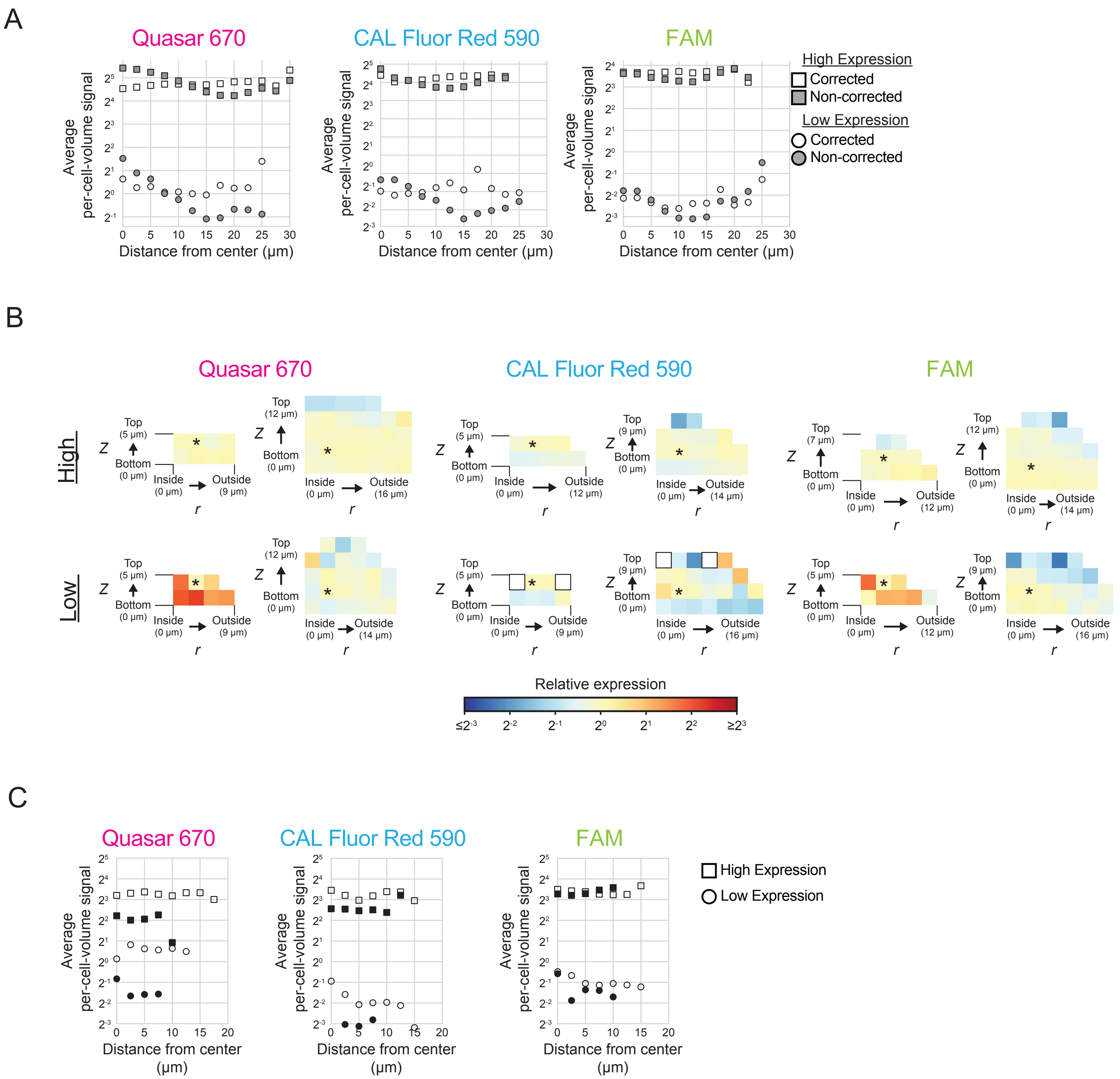

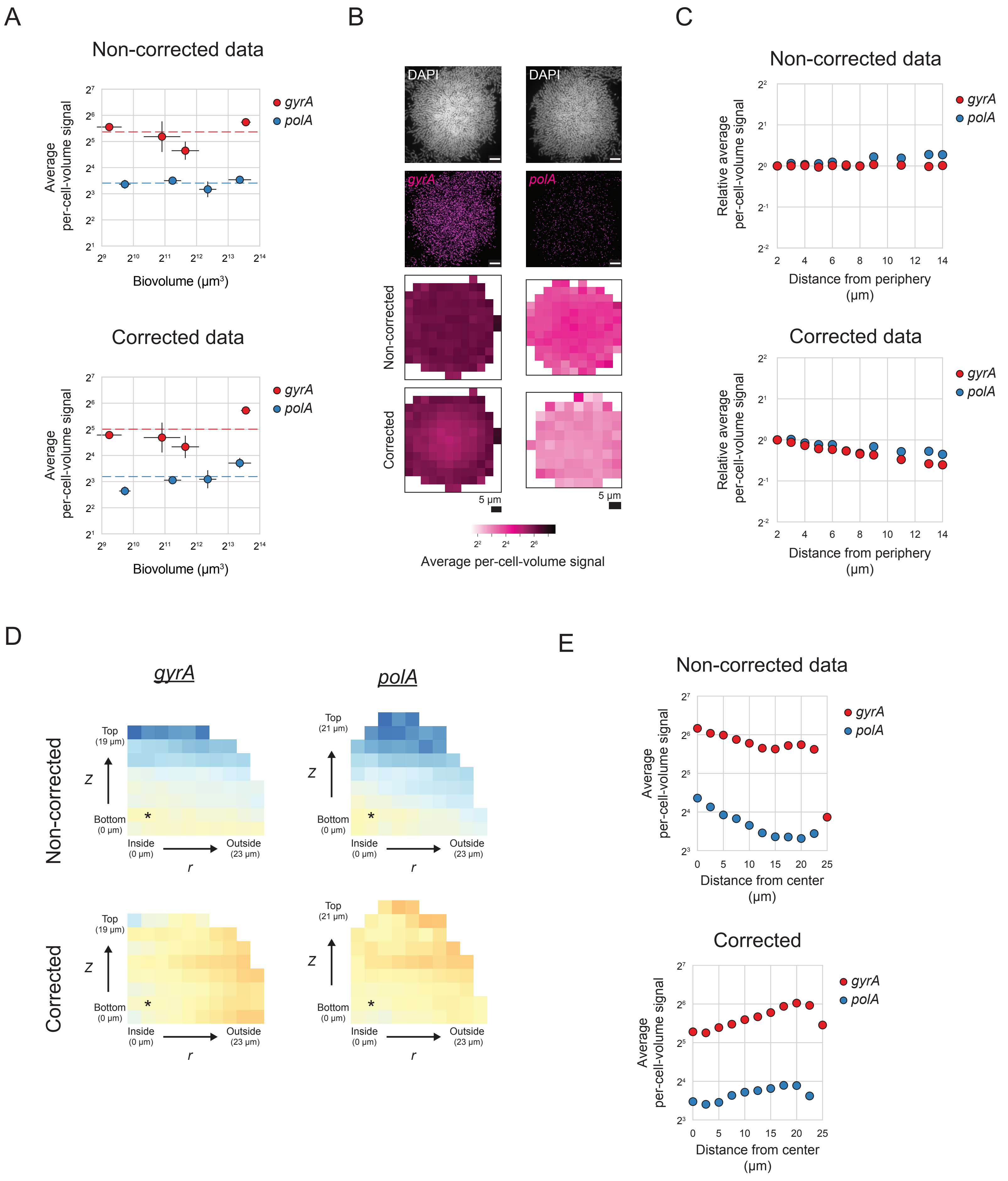

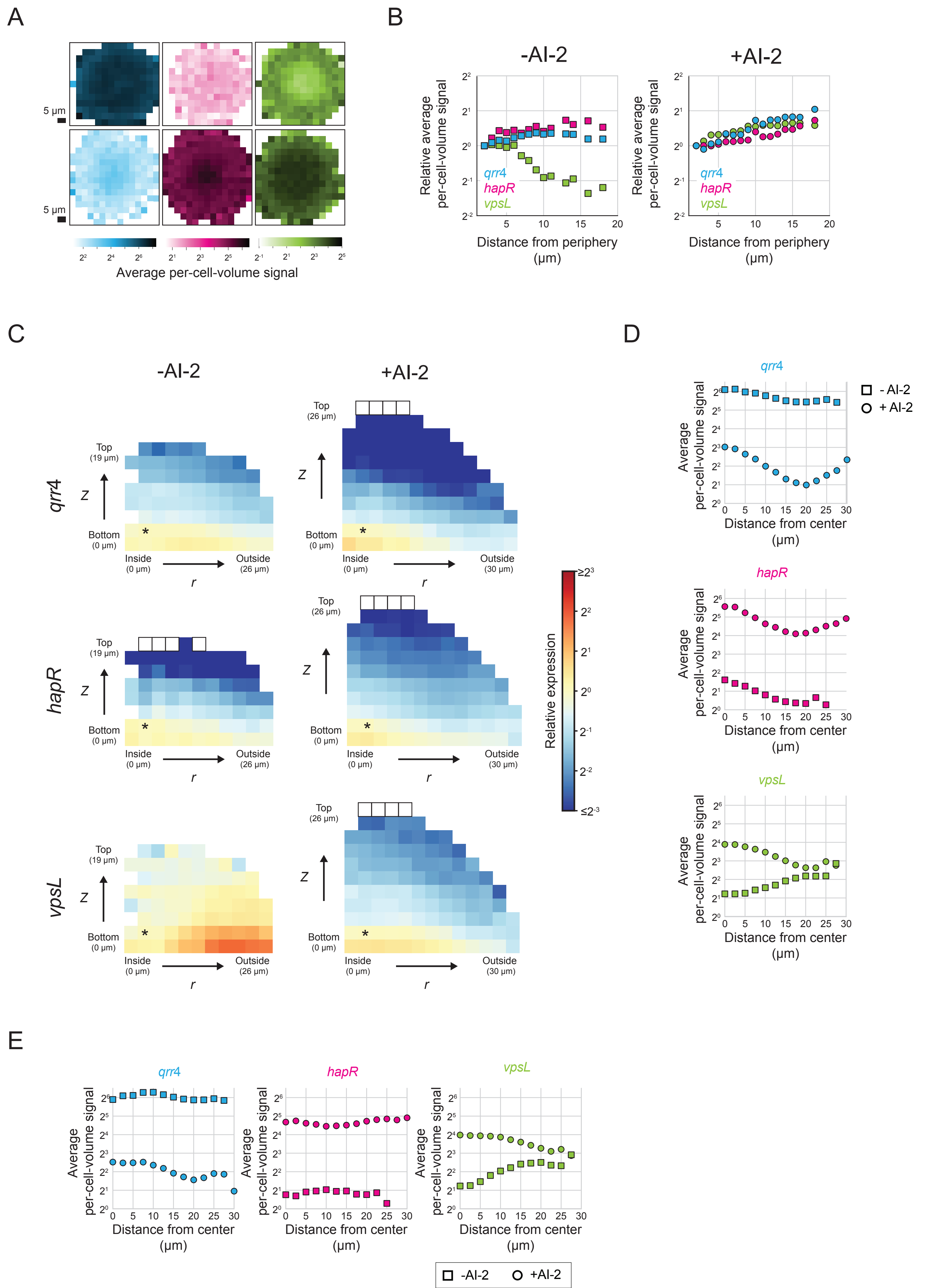

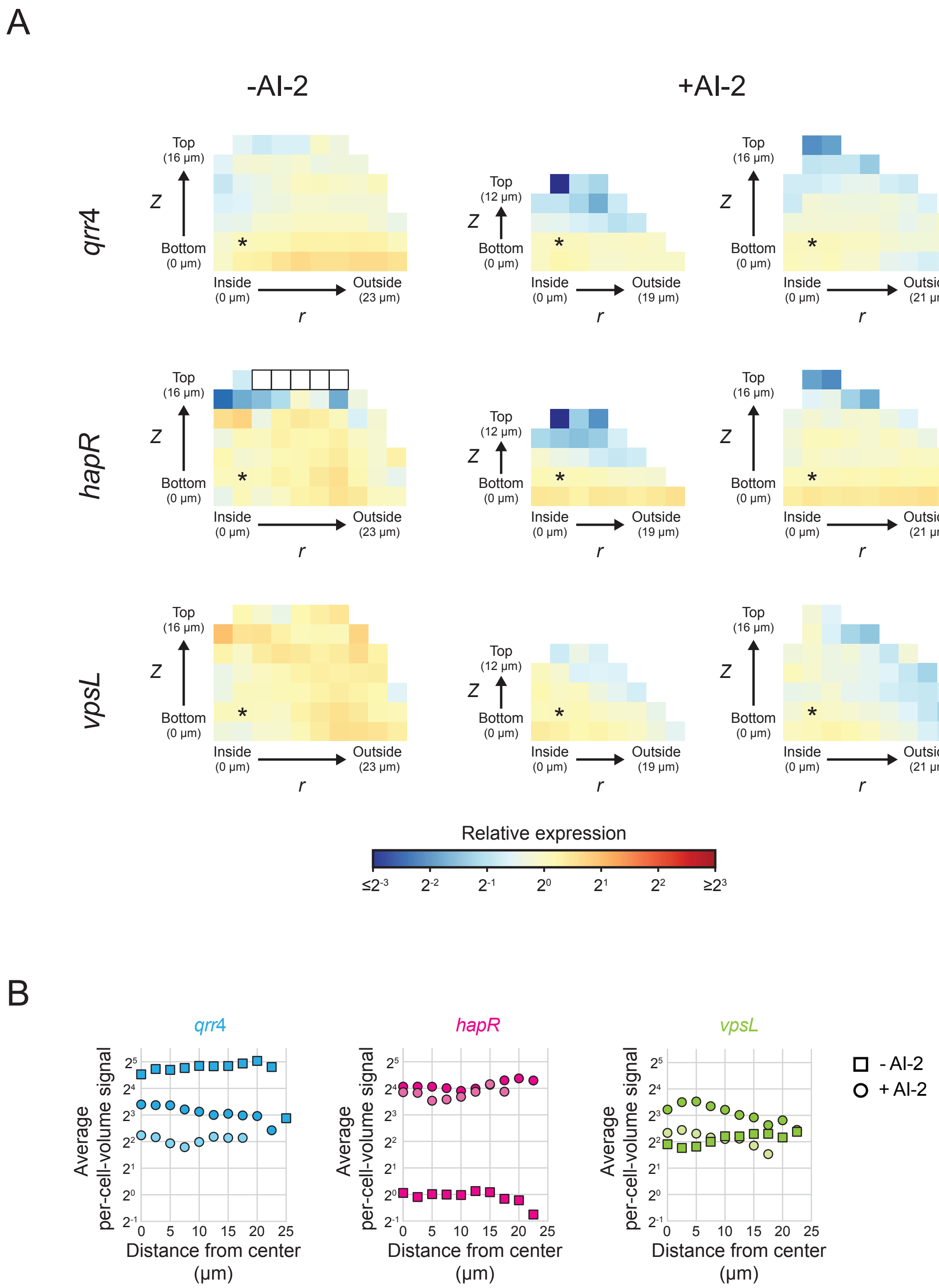

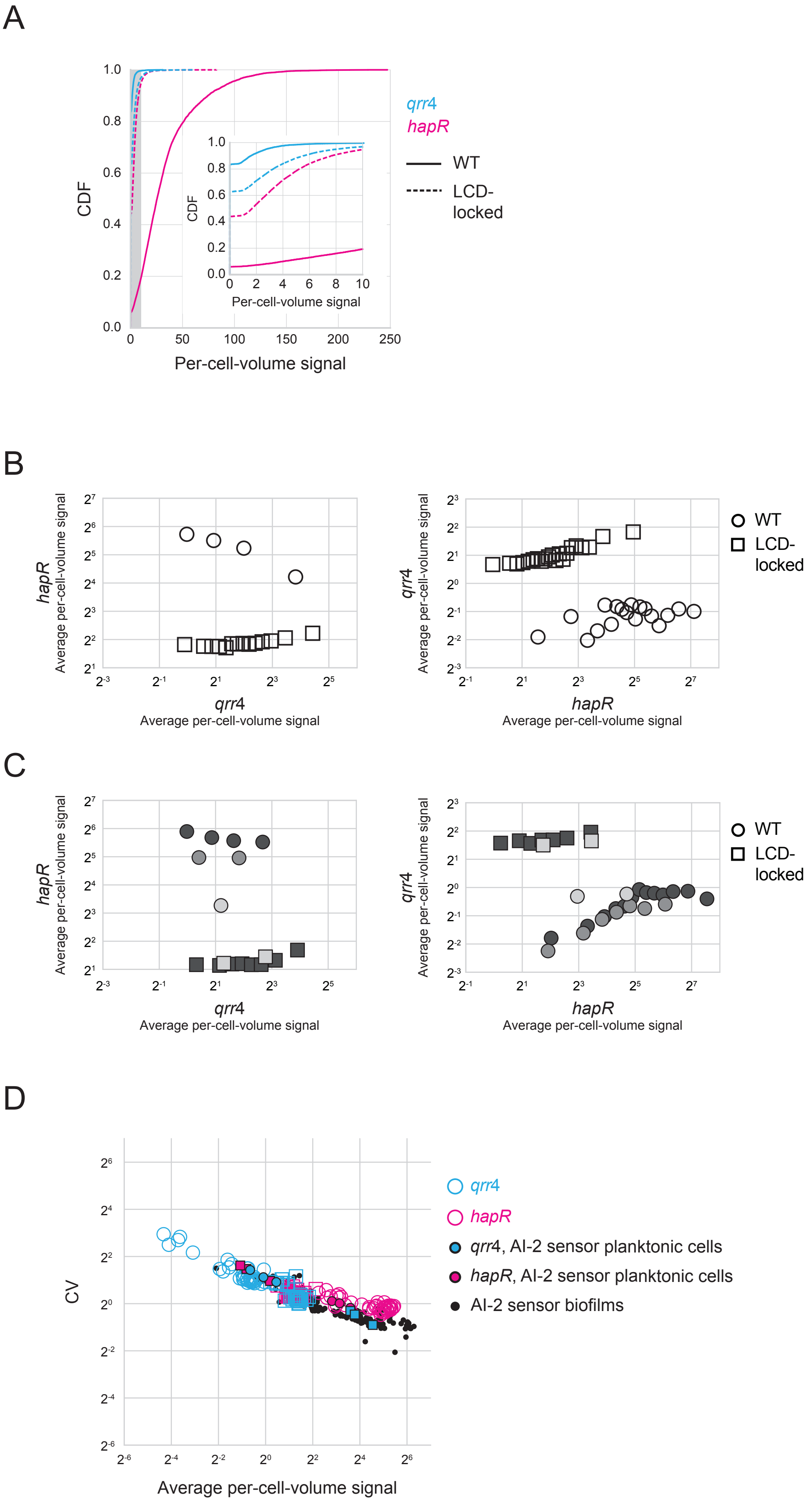

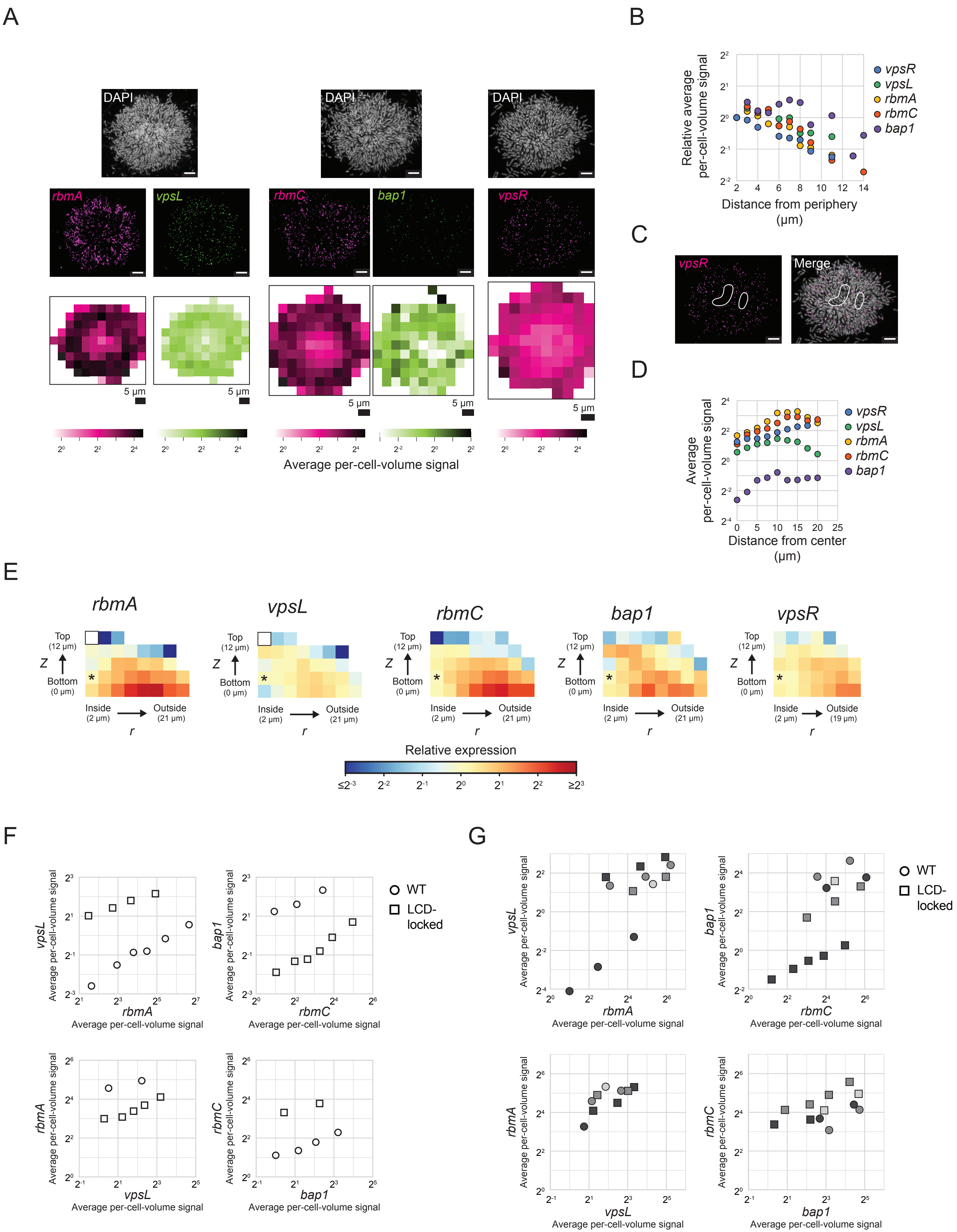
