## Supplementary material for "Cell-scale gene-expression measurements in *Vibrio cholerae* biofilms reveal spatiotemporal patterns underlying development": S1 Data

### S2A, non-corrected data

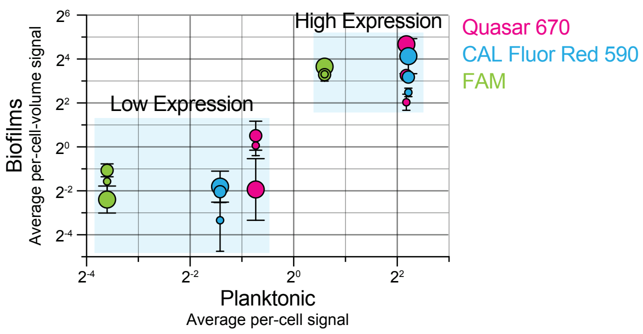


### S3B, non-corrected data

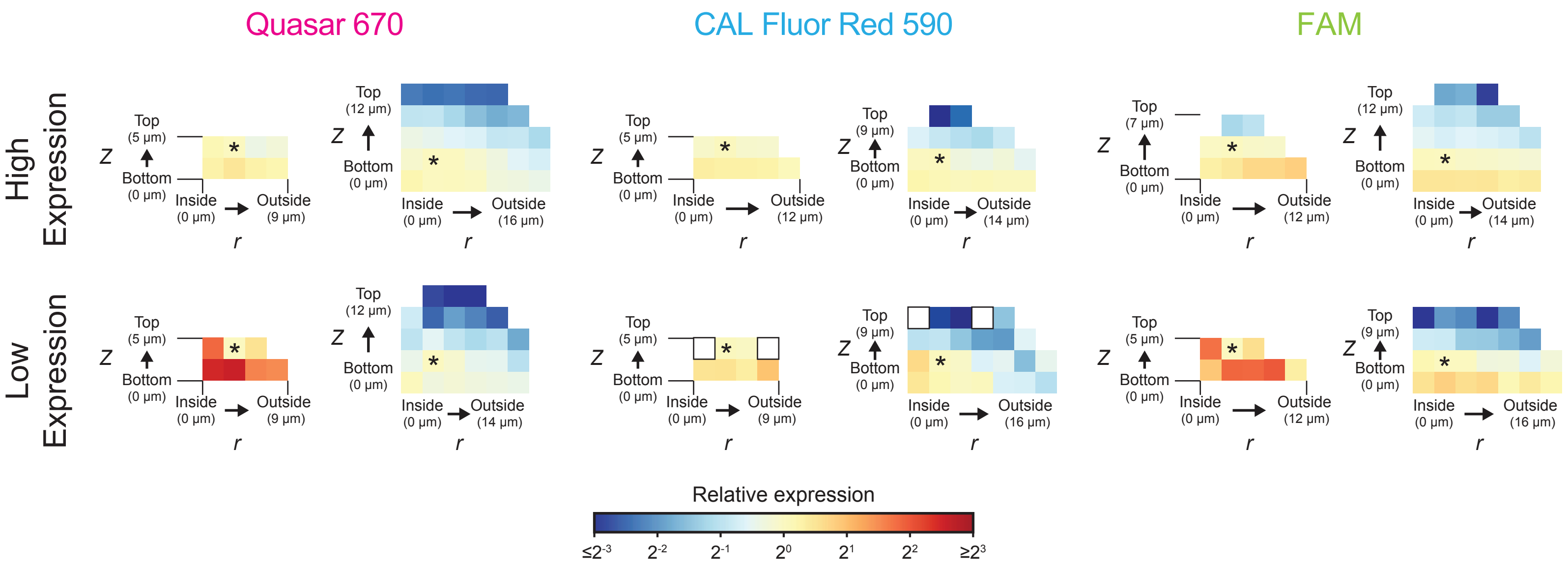


### S3C, non-corrected data

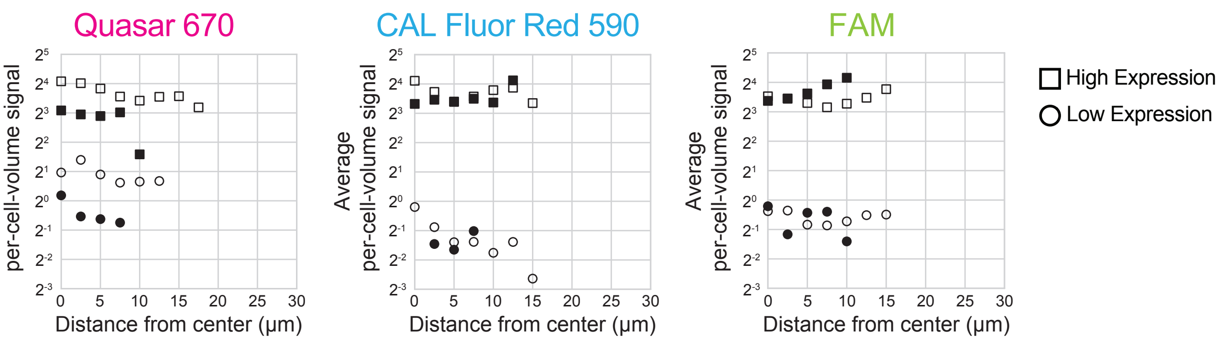

#### Fig 3A, non-corrected data

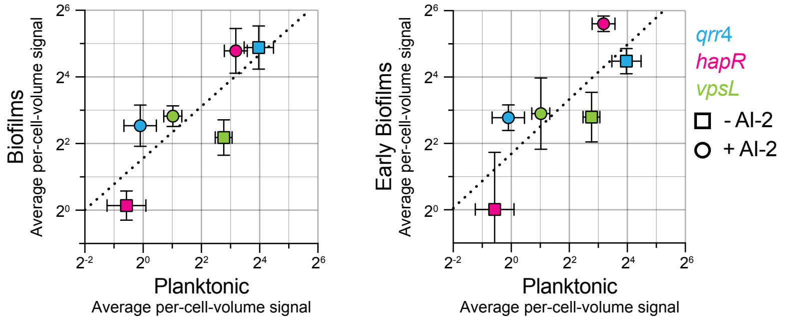


#### Fig 3B, non-corrected data

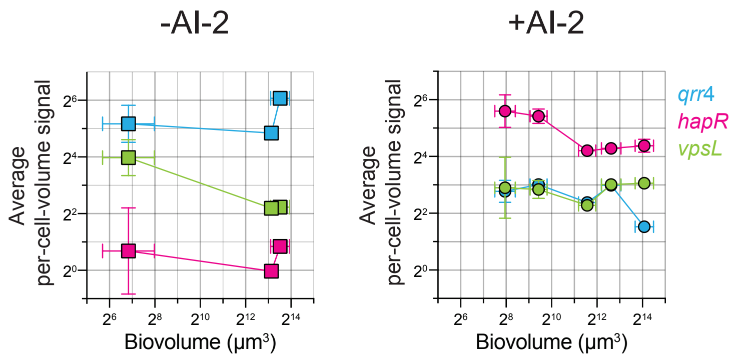


#### S6A, non-corrected data

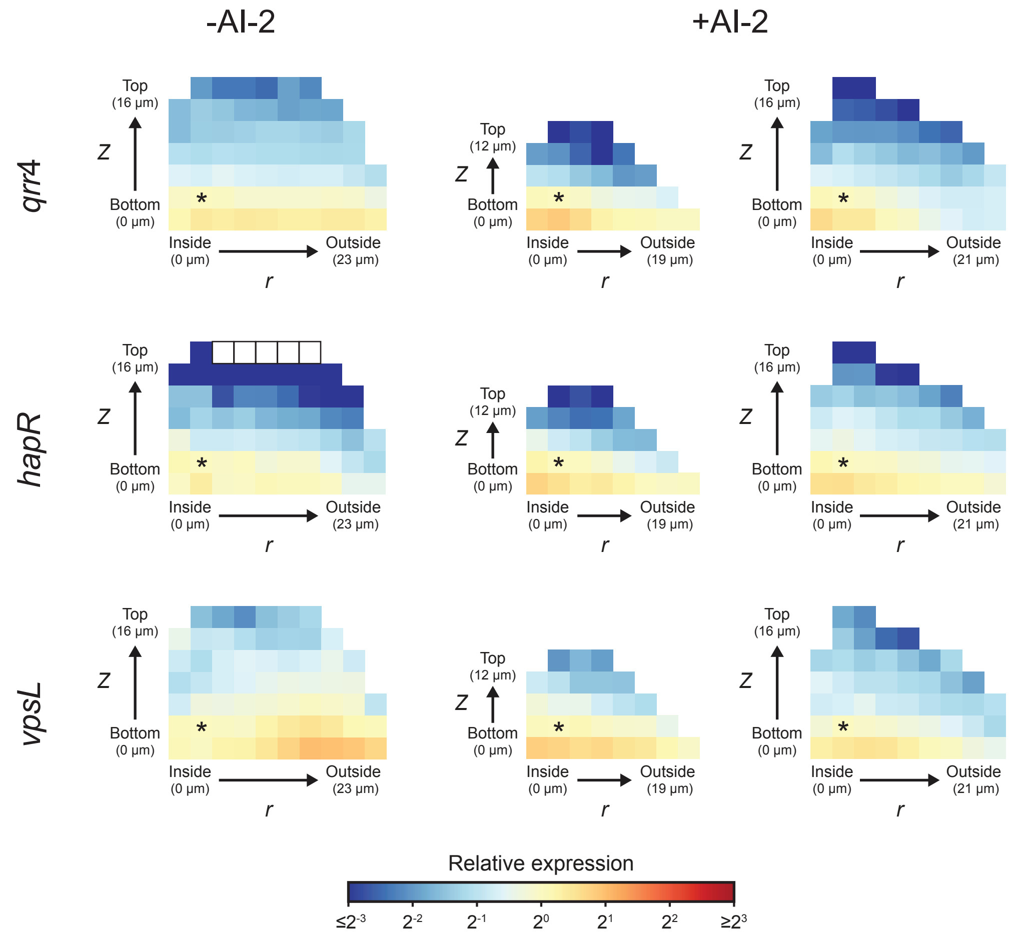


#### S6B, non-corrected data

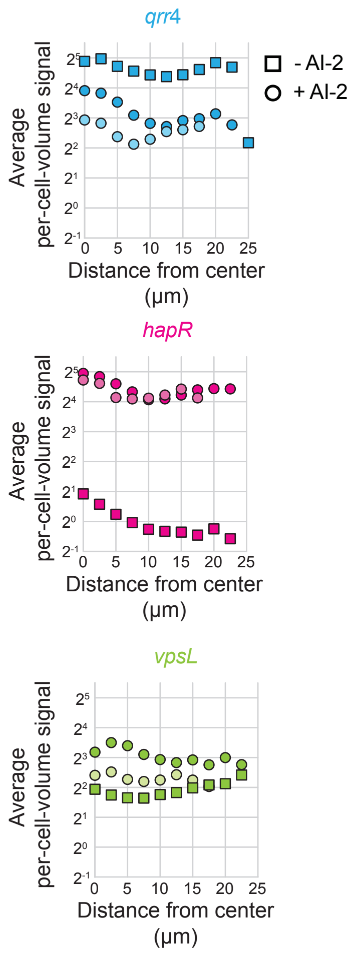

Fig 4A, non-corrected data

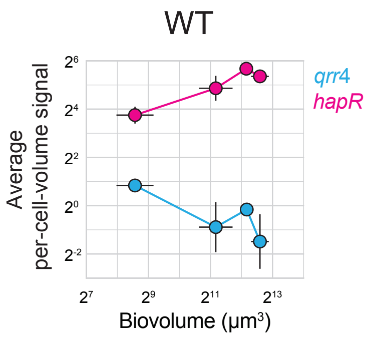


Fig 4B, non-corrected data

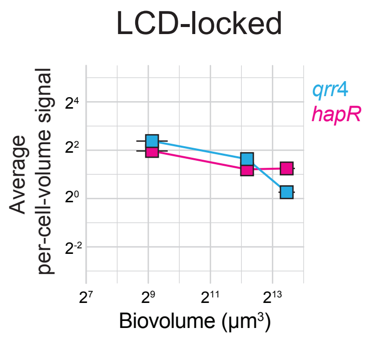


Fig 4C, non-corrected data

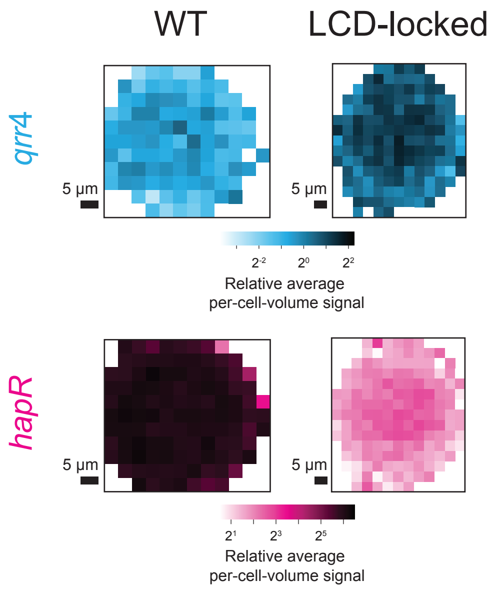


Fig 4D, non-corrected data

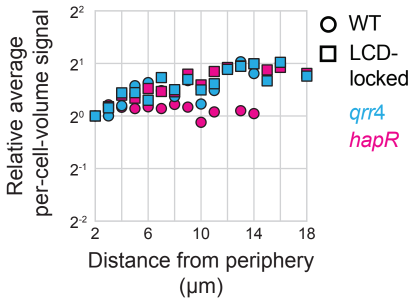


Fig 4E, non-corrected data

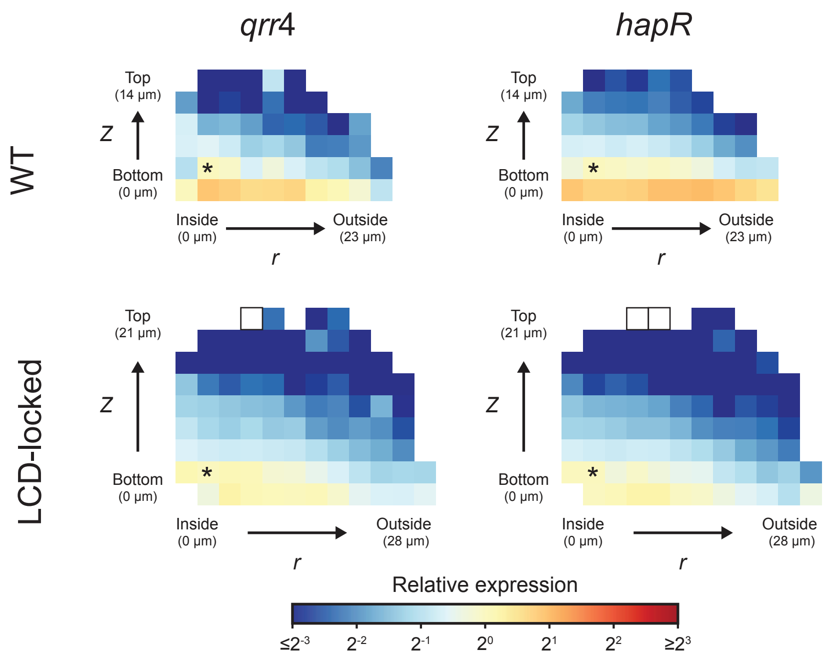


Fig 4F non-corrected data

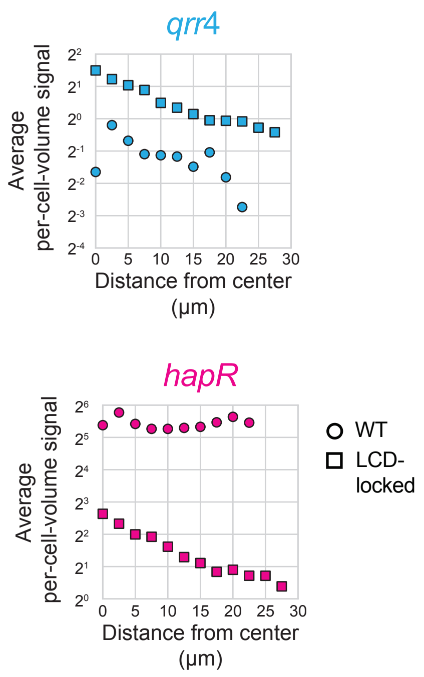

### Fig 5A, non-corrected data

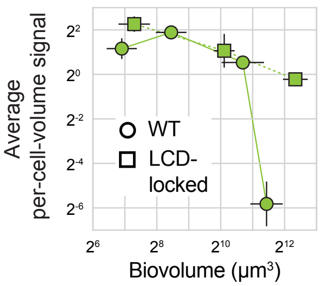


### Fig 5B, non-corrected data

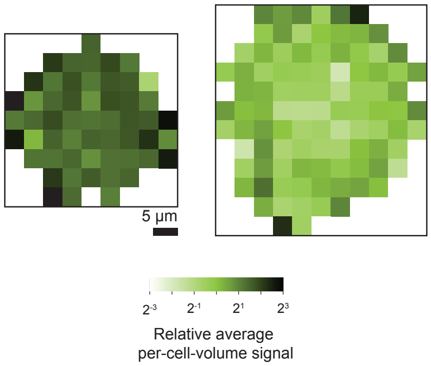


### Fig 5C, non-corrected data

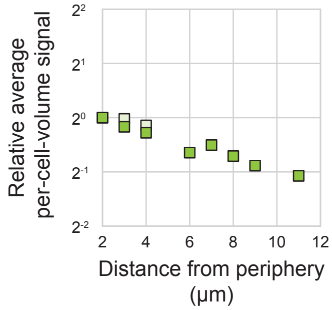


### Fig 5D, non-corrected data

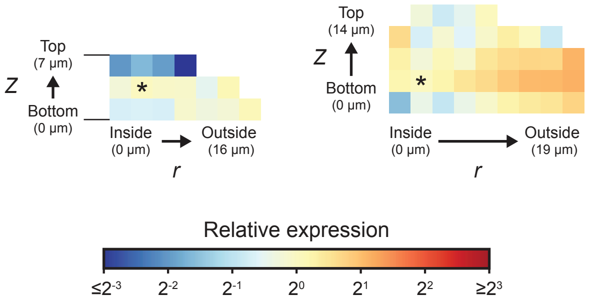


### Fig 5E, non-corrected data

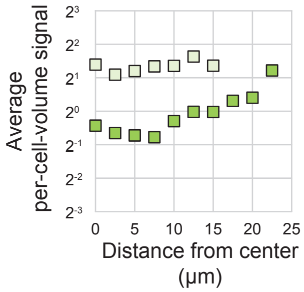

#### Fig 6A, non-corrected data

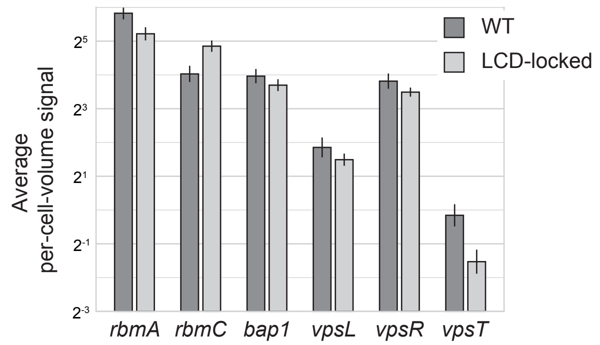


#### Fig 6B, non-corrected data

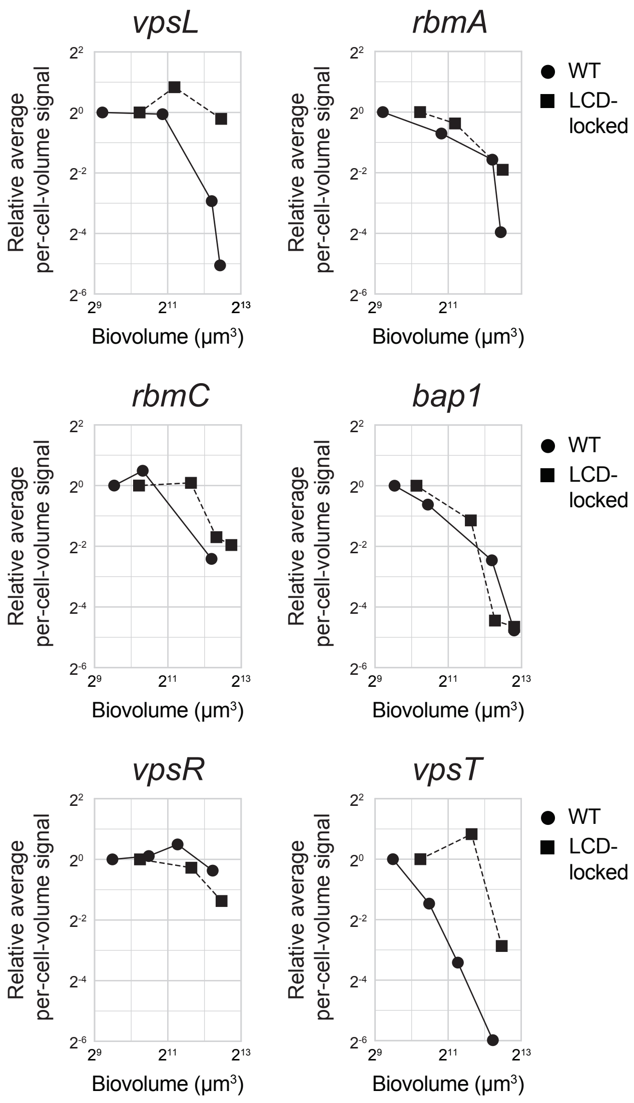


#### Fig 6C, non-corrected data

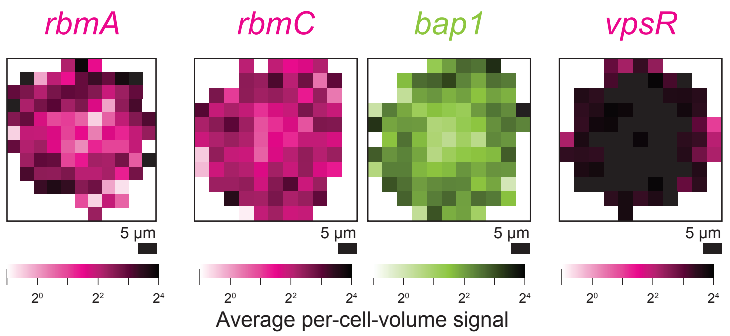


#### Fig 6D, non-corrected data

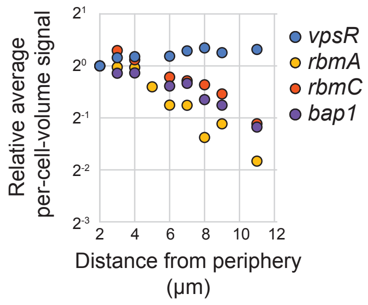


#### Fig 6E, non-corrected data

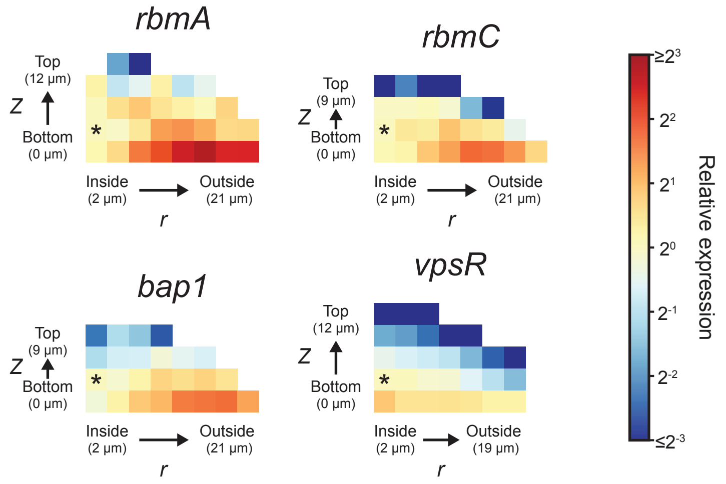


#### Fig 6F, non-corrected data

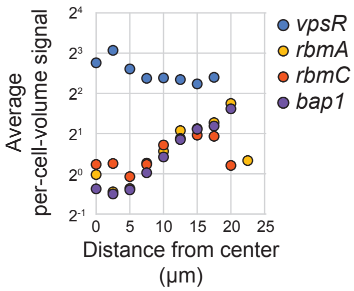

#### S8A, non-corrected data

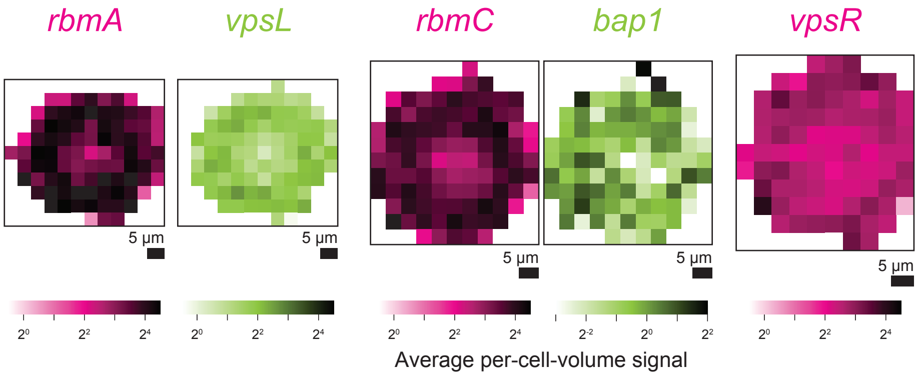


#### S8B, non-corrected data

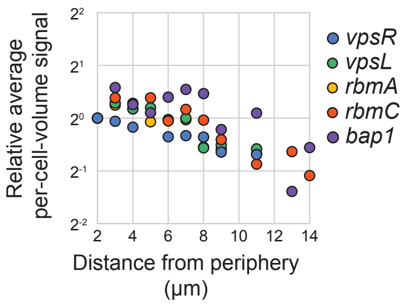


#### S8D, non-corrected data

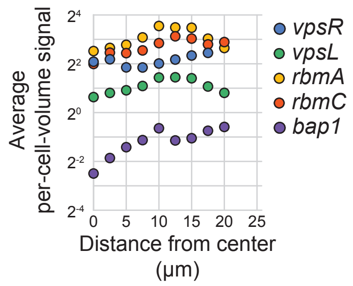


#### S8E, non-corrected data

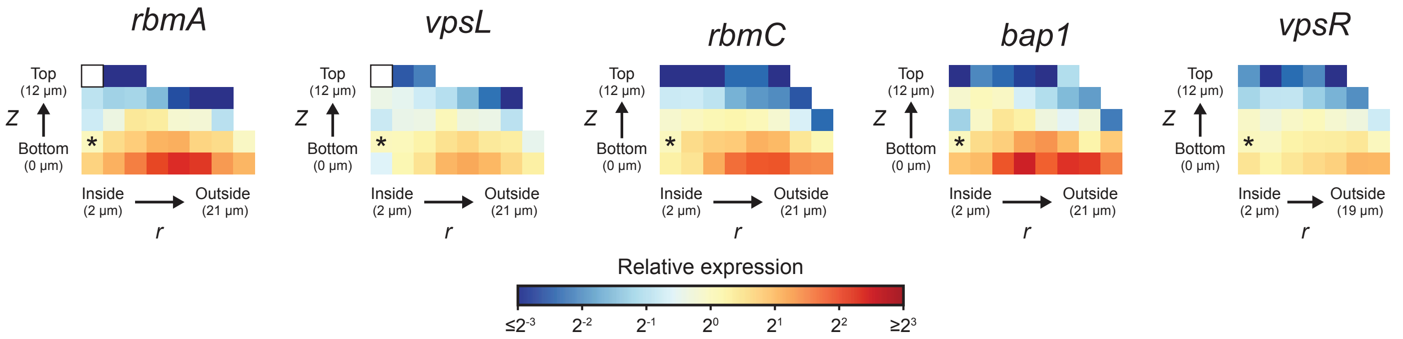
